# High emotional reactivity is associated with activation of a molecularly distinct hippocampal-amygdala circuit modulated by the glucocorticoid receptor

**DOI:** 10.1101/2022.06.01.494356

**Authors:** Qiang Wei, Vivek Kumar, Shannon Moore, Fei Li, Geoffrey G. Murphy, Stanley J. Watson, Huda Akil

**Affiliations:** Michigan Neuroscience Institute, University of Michigan, Ann Arbor, MI 48109.

## Abstract

Emotions are characterized not only by their valence but also by whether they are stable or labile. Yet, we do not understand the molecular or circuit mechanisms that control the dynamic nature of emotionality. We have shown that glucocorticoid receptor overexpression in the forebrain (GRov) leads to a highly reactive mouse with increased anxiety behavior coupled with greater swings in emotional responses. This phenotype is established early in development and persists into adulthood. However, the neural circuitry mediating this lifelong emotional lability remains unknown. In the present study, optogenetic stimulation in ventral dentate gyrus (vDG) of GRov mice led to a greater range and a prolonged duration of anxiety behavior. cFos expression showed that the amplified behavioral response to vDG activation in GRov mice is coupled to increased neuronal activity in specific brain regions. Relative to wild type mice, GRov mice displayed glutamatergic/GABAergic activation imbalance in ventral CA1 (vCA1) and selectively increased glutamatergic activation in the basal posterior amygdaloid complex. Fluorescence *in situ* hybridization chain reaction studies showed that forebrain GR overexpression led to increased activation of molecularly distinct subpopulations of neurons within the hippocampus and the posterior basolateral amygdala (pBLA). Increased cFos labeling was observed in the Calbindin 1 (Calbn1^+^) glutamatergic neurons in vCA1 and in the DARPP-32/Ppp1r1b^+^ glutamatergic neurons in pBLA. We propose that a molecularly distinct hippocampal-amygdala circuit is shaped by stress early in life and tunes the dynamics of emotional responses.

## Introduction

Mood and affective disorders are typically characterized by altered emotion regulation. Emotions may be more intense — e.g., highly positive or negative. But they can also exhibit different dynamics (Koenigsberg, 2010; Renaud and Zacchia, 2012). For example, in clinical depression or anxiety disorders, a feature of the negative affect is its persistence even in the face of a less stressful context (Bowen et al., 2004, 2011). By contrast, some psychiatric conditions such as bipolar disorder and borderline personality disorder are characterized by increased affective lability, whereby emotions appear unstable, easily changeable in the absence of a clear or significant external trigger (Henry et al., 2001; Marwaha et al., 2014; Taylor et al., 2021). The neural circuitry associated with the stability or lability of emotional responses is not well understood.

The hippocampus (HC) has long been implicated in learning and memory, but also has an important role in the regulation of mood (Femenía et al., 2012; Kim and Diamond, 2002). Recent optogenetic studies have corroborated the previously recognized critical role of HC in the processing of spatial memory (Buzsáki and Moser, 2013) as well as nonspatial memory which includes valence-related contextual information, e.g., contextual fear memory (Felix-Ortiz and Tye, 2014; Goosens, 2011). The distinct functions of HC arise from the anatomical and genetic heterogeneity along the dorsal-ventral axis with dorsal HC (dHC) specializing in accurate spatial navigation and episodic memory processes (Mably et al., 2017; O’Keefe, 1979) and ventral HC (vHC) specializing in affective and motivated behaviors (Ciocchi et al., 2015; Fanselow and Dong, 2010; Strange et al., 2014; Kheirbek and Hen, 2011). This functional specialization is the result of distinct efferent projections. dHC projects to regions such as entorhinal cortex, retrosplenial cortex, and septum known to mediate context-associations and spatial memory processes (Canteras and Swanson, 1992; Cenquizca and Swanson, 2007), while vHC targets limbic structures such as medial prefrontal cortex (mPFC), amygdala, nucleus accumbens (NAcc), and hypothalamus to directly mediate anxiety and mood-related behaviors (Canteras and Swanson, 1992; Jay and Witter, 1991; Jimenez et al., 2018; Kishi et al., 2006; Phillipson and Griffiths, 1985; Tovote et al., 2015). Recent studies that have modulated either hippocampal efferents, such as vHC to lateral septum and mPFC (Padilla-Coreano et al., 2016; Parfitt et al., 2017) or hippocampal afferents such as basolateral amygdala (BLA) to vHC inputs (Felix-Ortiz et al., 2013) have further supported the role of vHC in the regulation of emotions. However, a potential role of the hippocampus in regulating the dynamics of emotional reactivity remains relatively unexplored.

One clear link between the hippocampus and emotionality is the role of this structure in controlling the stress response. The HC expresses the highest levels of GR, an organizational molecule that has been implicated in both the perception and the termination of the stress response (McEwen and Akil, 2000). In order to understand the role of brain GR in emotions, we created a forebrain-specific GR overexpressing mouse (GRov) driven by the CaMKIIa promoter. The GRov mouse exhibits a very distinct phenotype characterized by enhanced responsiveness to both negative and positive emotional stimuli, including the behavioral impact of antidepressants and sensitization to psychoactive drugs (Wei et al., 2004, 2012). We also created an inducible GRov mouse to study developmental mechanisms and showed that GR overexpression prior to weaning was necessary and sufficient to induce the entire phenotype. Gene expression profiling showed that both lifelong and early life GR overexpression resulted in significant changes in hippocampal glutamatergic and calcium signaling, some of which are implicated in bipolar disorder (Wei et al., 2007, 2012). Indeed, many features of GRov mice are consistent with the clinical features of hypomania and mania, and it is notable that these states can be induced during acute corticosteroid therapy (Brown, 2009; Kenna et al., 2011). Thus, the GRov mouse represents a useful model to investigate the neural mechanisms of emotional dynamics that are dysregulated in certain mood disorders.

Given the canonical trisynaptic loop within HC from the input node of the DG to CA3 and to the output node CA1 (Treves and Rolls, 1994), the DG serves as a logical entry-point for the pattern separation of cortical inputs during memory encoding and downstream emotional processing (Leutgeb et al., 2007).

Optogenetic stimulation of granule cells in the vDG has been shown to cause decreased anxiety and increased risk-taking behavior (Kheirbek et al., 2013). In the present study, we asked whether GR in the DG modulates the magnitude and duration of emotional responses and sought to uncover the associated neural circuitry. We compared the impact of optogenetic stimulation of either vDG or dDG on anxiety responses in GRov mice relative to their wild type littermates. We coupled the stimulation and behavioral findings with anatomical analyses to evaluate the extent of cellular activation throughout the brain as measured by cfos, and then characterized the molecular nature of the two most differentially activated brain regions. Our findings reveal how a stress gene can retune a neural circuit that can impact emotional reactivity.

## Results

### Differential Emotional Reactivity to the Activation of dDG and vDG in GRov Mice

We first examined the effects of modulating activity in the dDG or vDG on acute control of emotional reactivity in GRov mice. To manipulate neuronal activity of dDG or vDG, we microinjected AAV5-CaMKIIa-ChR2(H134R)-eYFP virus into dDG (Fig. 1*A*) or vDG (Fig. 2*A*) of adult GRov mice and WT littermates. Animals exhibited robust expression of opsin ChR2-eYFP in dDG (Fig. 1*B*) or vDG (Fig. 2*B*) after 3 weeks of the virus injection. For optogenetic stimulation in dDG, both GRov mice and WT littermates were implanted with fiber optics targeting the dDG and tested for 18 min in the elevated plus maze (EPM) with 6 min light off (Off1), 6 min light on (On), 6 min light off (Off2) epochs (Fig. 1*A*).

**Fig. 1.**
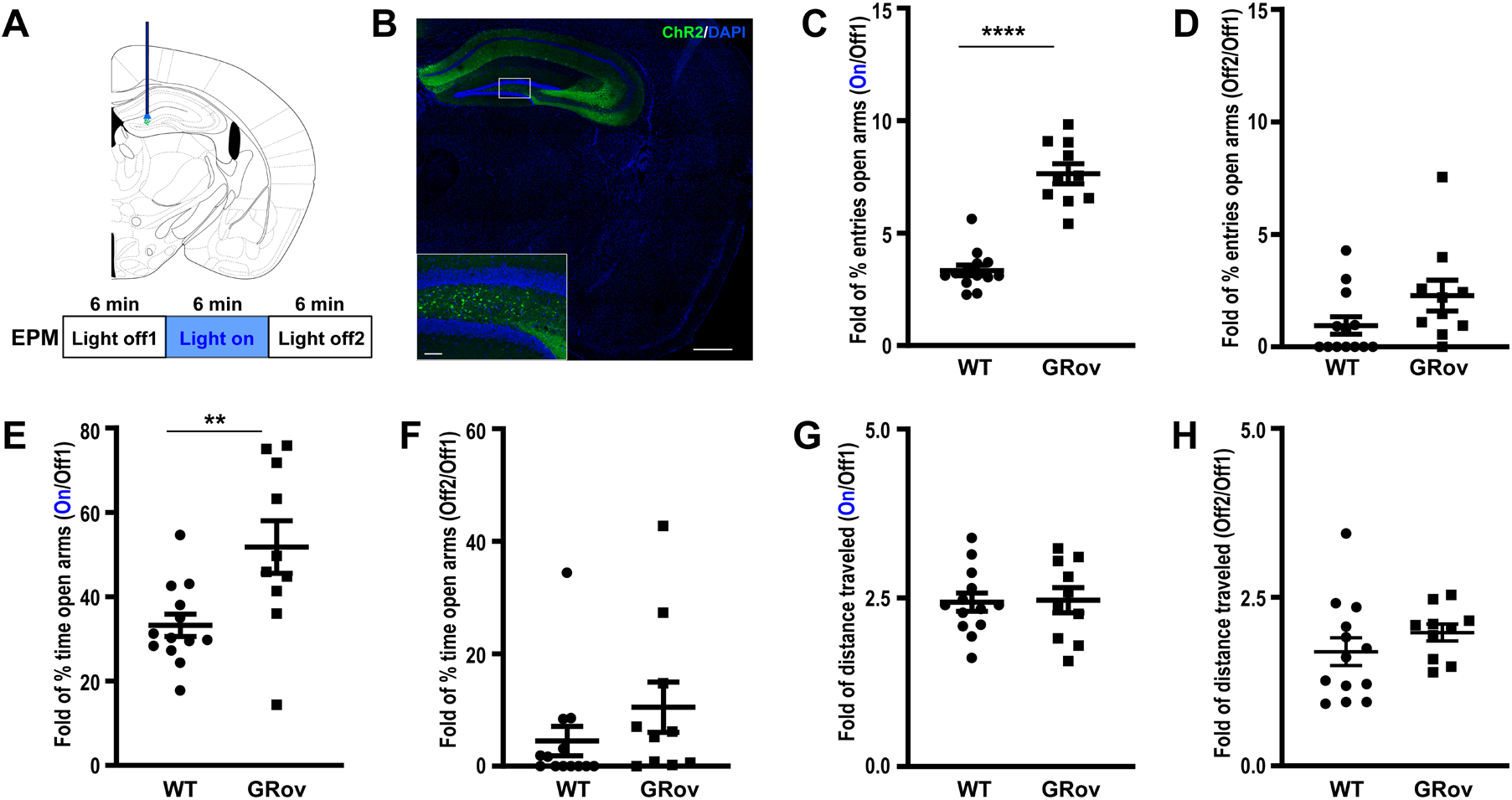
Increased emotional response to the optogenetic activation in dDG of GRov mice. (*A*) Experimental design. Mice expressing ChR2-eYFP were implanted in the dDG then tested for 18 min in the EPM with 6 min blue light off (Off1), 6 min blue light on (On), 6 min blue light off (Off2) epochs. (*B*) Expression of ChR2-eYFP in dDG. (*C*) Following optogenetic stimulation in dDG, GRov mice exhibited a greater response range in entering into open arms than WT. Unpaired two tailed *t* test: *t*(21) = 9.04, \*\*\*\**P* < 0.0001. (*D*) There is no genotype difference in response range of entering into open arms after the termination of light between GRov and WT mice. Unpaired two tailed *t* test: *t*(21) = 1.77, *P* =0.09. (*E*) Following optogenetic stimulation in dDG, GRov mice exhibited a greater response range in time spent in open arms than WT. Unpaired two tailed *t* test: *t*(21) = 3, \*\**P* < 0.01. (*F*) There is no genotype difference in response range of time spent in open arms after the termination of blue light between GRov and WT mice. Unpaired two tailed *t* test: *t*(21) = 1.22, *P* =0.24. (*G* and *H*) There is no genotype difference in response range in distance traveled in EPM following the photostimulation or after the termination of blue laser stimulation in dDG. Unpaired two tailed *t* test: *t*(21) = 0.15, *P* =0.89, for photostimulation epoch; *t*(21) = 1.1, *P* =0.26, for blue laser off epoch after the stimulation. GRov, *n* = 10 mice; WT, *n* = 13 mice.

**Fig. 2.**
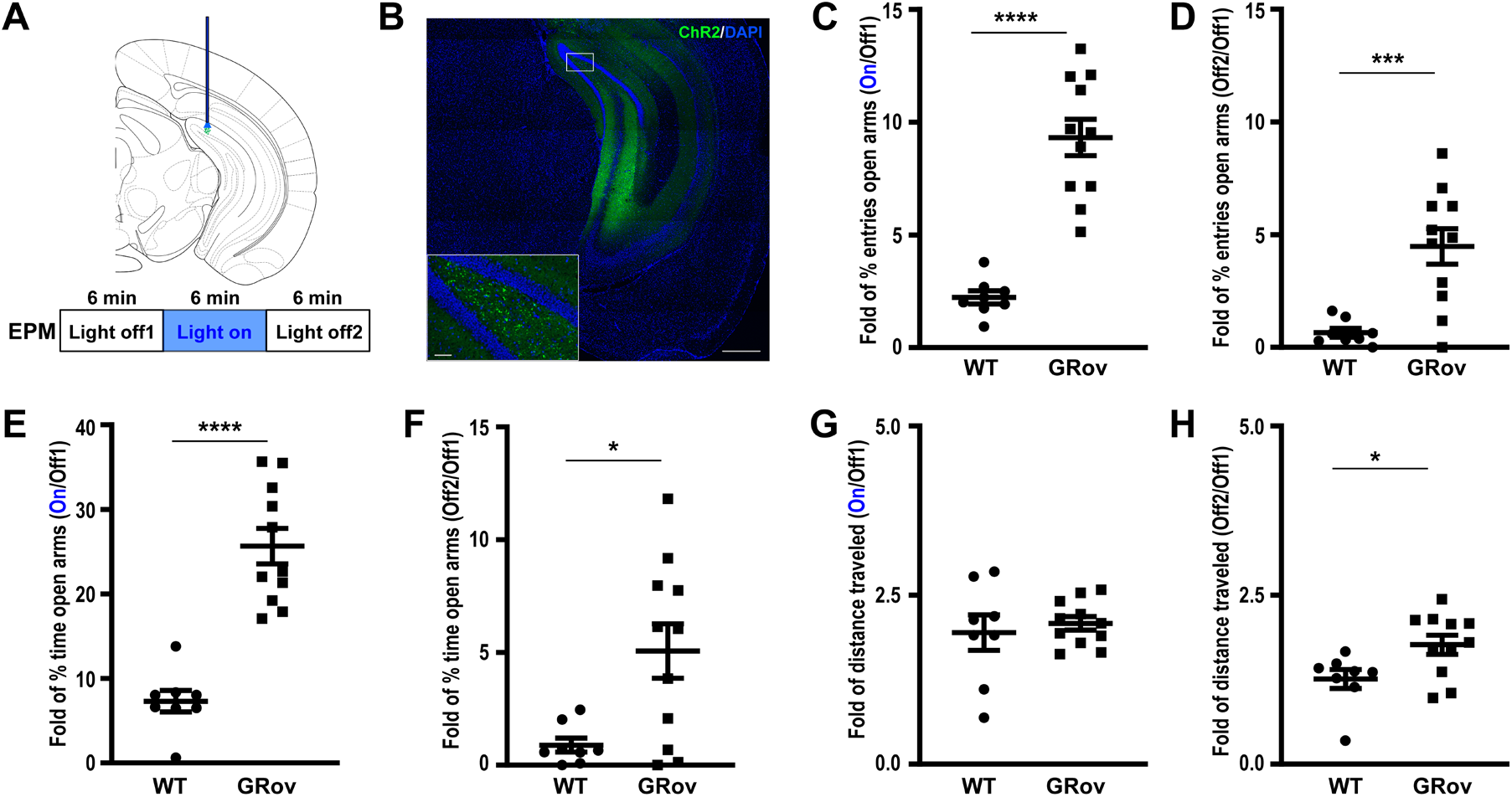
Increased and prolonged emotional reactivity to the optogenetic stimulation in vDG of GRov mice. (*A*) Experimental design. Mice expressing ChR2-eYFP were implanted in the vDG then tested for 18 min in the EPM with 6 min blue light off (Off1), 6 min blue light on (On), 6 min blue light off (Off2) epochs. (*B*) Expression of ChR2-eYFP in vDG. (*C*) Following optogenetic stimulation in vDG, GRov mice exhibited a greater response range in entering into open arms than WT. Unpaired two tailed *t* test: *t*(17) = 7.17, \*\*\*\**P* < 0.0001. (*D*) Moreover, GRov mice continuously displayed a greater response range in entering open arms after the termination of light stimulation than WT. Unpaired two tailed *t* test: *t*(17) = 4.02, \*\*\**P* < 0.001. (*E*) Following optogenetic stimulation in vDG, GRov mice exhibited a greater response range in time spent in open arms than WT. Unpaired two tailed *t* test: *t*(17) = 6.77, \*\*\*\**P* < 0.0001. (*F*) GRov mice extended their greater response range in time spent in open arms after the termination of light stimulation than WT mice. Unpaired two tailed *t* test: *t*(17) = 2.88, \**P* < 0.05. (*G*) There is no genotype difference in response range in distance traveled in EPM following the blue laser stimulation. Unpaired two tailed *t* test: *t*(17) = 0.54, *P* = 0.6. (*H*) GRov mice displayed a greater response range in distance traveled in EPM after the termination of light stimulation. Unpaired two tailed *t* test: *t*(17) = 2.46, \**P* < 0.05. GRov, *n* = 11 mice; WT, *n* = 8 mice.

#### Dorsal Dentate Gyrus Activation

Optogenetic activation of dDG induced robust and reversible increases in entries into the open arms of the EPM as well as the time spent in the open arms for both GRov and WT mice (*SI Appendix*, Fig. S1 A and B). This was accompanied by increased total exploration (*SI Appendix*, Fig. S1C). Interestingly, GRov mice showed greater response in magnitude to the light stimulation in dDG than WT mice when compared to their respective pre-stimulation baselines (Fig. 1 *C* and *E*). There is no genotype difference in response range of entering into open arms or time spent in open arms of the EPM after the termination of light stimulation (Fig. 1 *D* and *F*). While optogenetic activation of dDG increased total distance traveled in EPM (*SI Appendix*, Fig. S1C), there is no genotype difference in response range in distance traveled in EPM following the photostimulation (Fig. 1*G*) or after the termination of light stimulation (Fig. 1*H*). Blue light illumination of the dDG in mice with control virus microinjection (AAV5- CaMKIIa-mCherry) in dDG did not induce any increase in entries into the open arms or time spent in the open arms of the EPM (*SI Appendix*, Fig. S2 A and B). Mice with control virus injection showed normal habituation behavior in the EPM across testing epochs (*SI Appendix*, Fig. S2 A and B) with comparable general exploration (*SI Appendix*, Fig. S2C). Thus, the observed behavioral phenotype in opsin ChR2-eYFP-expressing mice was specific to the activation of dDG. These results suggest that GRov mice had a greater range in their response to the activation of dDG.

#### Ventral Dentate Gyrus Activation

For optogenetic activation in vDG, both GRov mice and WT littermates were implanted with fiber optics targeting the vDG and tested for 18 min in EPM with 6 min light off (Off1), 6 min light on (On), 6 min light off (Off2) epochs (Fig. 2*A*). Optogenetic activation of vDG induced robust increases in entries into the open arms and in time spent in the open arms of the EPM in both GRov and WT mice (*SI Appendix*, Fig. S3 A and B), accompanied by increased total exploration (*SI Appendix*, Fig. S3C). GRov mice showed increased response magnitude during the vDG stimulation (Fig. 2 *C* and *E*) and this phenotype extended even after the termination of the light stimulation (Fig. 2 *D* and *F*), indicating greater and more prolonged emotional reactivity in GRov animals. While optogenetic activation of vDG increased general exploration in EPM (*SI Appendix*, Fig. S3C), there is no genotype difference in response range in distance traveled in EPM following the photostimulation (Fig. 2*G*). However, GRov mice showed increased general exploration (*SI Appendix*, Fig. S3C) after the termination of light stimulation and had a greater response range in distance traveled in EPM (Fig. 2*H*) relative to WT. These results demonstrate that GRov mice displayed a greater range in their response to the light stimulus and a prolonged effect on anxiety behavior following the activation of vDG. Thus, the optogenetic activation in vDG rapidly shifted the nature and duration of affective state in GRov mice from more anxious to less anxious than WT, indicating GRov mice are more prone to shift from one emotional state to another.

### Distinct Neuronal Activation Patterns Following dDG and vDG Activation in GRov Mice

We then evaluated the neuronal activation pattern across the brain using cFos mRNA expression as an activity marker.

#### Dorsal Dentate Gyrus

Optogenetic stimulation in the dDG led to increased cFos expression in a limited set of brain regions in both GRov and WT mice (Fig. 3*A*). These included the cerebral cortex (PFC, prelimbic, infralimbic and cingulate cortex), primary/secondary motor and somatosensory cortex (M1/2 and S1), lateral septum (LS), bed nucleus of the stria terminalis (BNST), thalamus, paraventricular hypothalamic nucleus (PVN), hippocampus (CA1, CA2, CA3, and DG), hypothalamic nuclei, and amygdaloid complex (Fig. 3*B* and *SI Appendix*, Table S1). Most of these activated regions are associated with the modulation of emotional responses. Moreover, an intense cFos response was observed in DG, CA1, CA3, and LS following dDG stimulation (Fig. 3*B* and *SI Appendix*, Table S2). Next, we compared the cFos response between GRov and WT mice following the dDG stimulation. We found that stimulation led to a significant increase in cFos expression in LS, DG, CA1, and pBLA of GRov mice as compared to WT (Fig. 3*B* and *SI Appendix*, Table S2), indicating a higher neuronal activity in the respective brain areas of GRov animals.

**Fig. 3.**
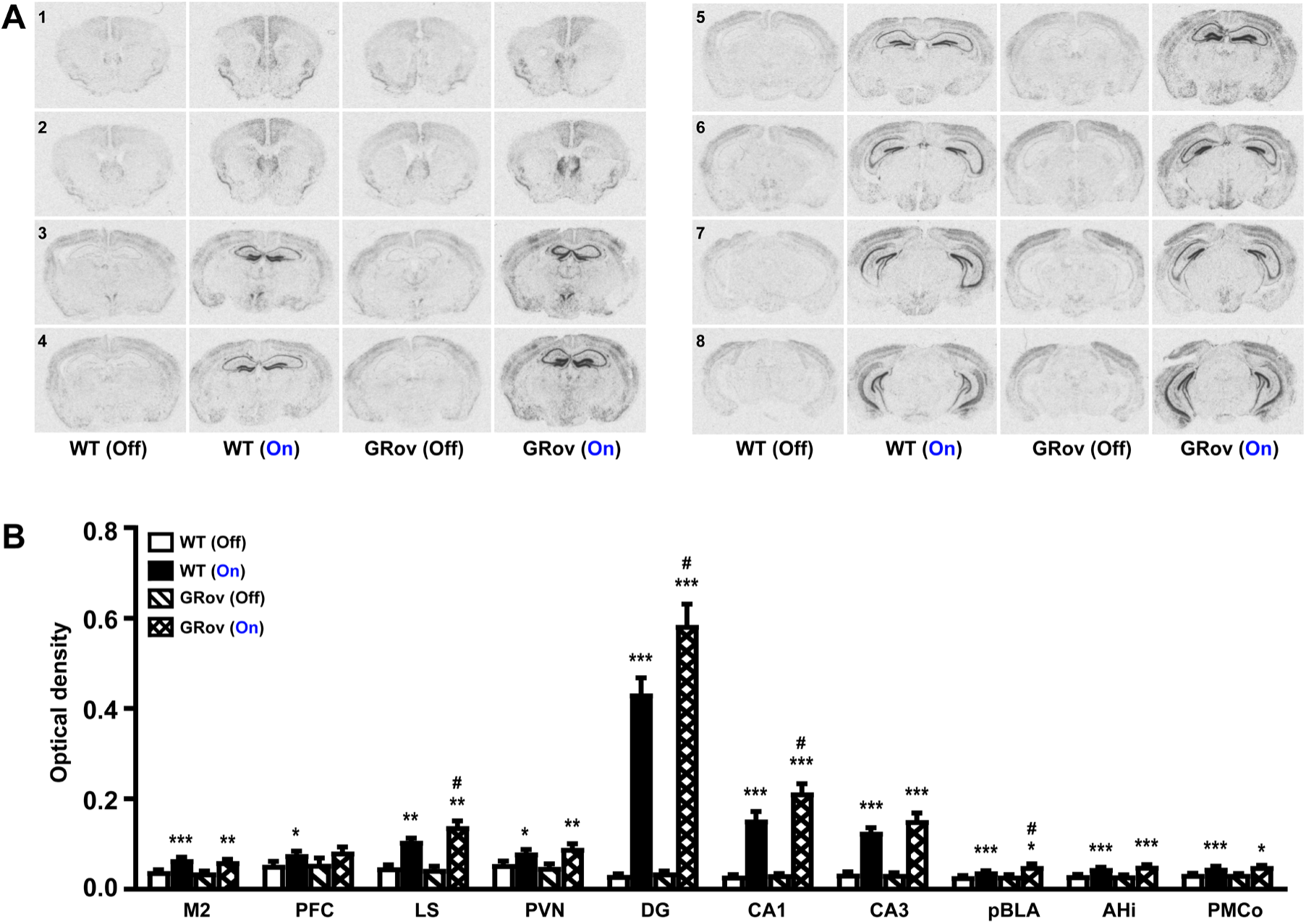
Neuronal activation following optogenetic stimulation in dDG of GRov mice. (*A*) The extent of neuronal activation in the brain following light stimulation in dDG as revealed by *In situ* hybridization (ISH). Higher cFos mRNA expression can be seen in key brain regions following activation of dDG compared to no stimulation group. Representative images are shown ranging from Bregma 1.18 mm (indicated by the no. 1) through Bregma -3.08 mm (indicated by the no. 8). (*B*) Images were quantitatively analyzed. All listed regions showed increased cFos expression as a result of stimulation in dDG. dDG stimulation led to a greater extent of cFos expression in LS, DG, CA1, and pBLA of GRov mice as compared to WT. Mean ± SEM. Unpaired two-tailed *t*-test: \**P* < 0.05, \*\**P* < 0.01, ***P < 0.001 versus respective no stimulation group; ^#^*P* < 0.05 versus WT group under the same condition; WT (light off), *n* = 5 mice; WT (light on), *n* = 7 mice; GRov (light off), *n* = 4 mice; GRov (light on), *n* = 7 mice.

#### Ventral Dentate Gyrus

In contrast, optogenetic stimulation of vDG resulted in a widespread cFos response in both GRov and WT mice (Fig. 4*A*), including cerebral cortex (PFC, prelimbic, infralimbic and cingulate cortex), primary/secondary motor and somatosensory cortex (M1/2 and S1), nucleus accumbens, LS, BNST, thalamus, PVN, hippocampus (CA1, CA2, CA3, and DG), hypothalamic nuclei, mammillary nucleus, and amygdaloid complex in particular pBLA, posterior basomedial amygdaloid nucleus, amygdalohippocampal area (AHi), posteromedial cortical amygdaloid nucleus (PMCo), and posterolateral cortical amygdaloid nucleus (Fig. 4*B* and *SI Appendix*, Table S3). Similar to dDG, stimulation of vDG resulted in a greater cFos response in DG and CA1 of GRov mice as compared to WT (Fig. 4*B* and *SI Appendix*, Table S4). Furthermore, vDG- stimulation led to a robust cFos mRNA expression in the basal posterior amygdaloid complex of GRov mice, including pBLA, AHi, and PMCo (Fig. 4 *A* and *B* and *SI Appendix*, Table S4). Importantly, the cFos response in all three amygdaloid subregions was significantly higher in GRov compared to WT (Fig. 4*B* and *SI Appendix*, Table S4).

**Fig. 4.**
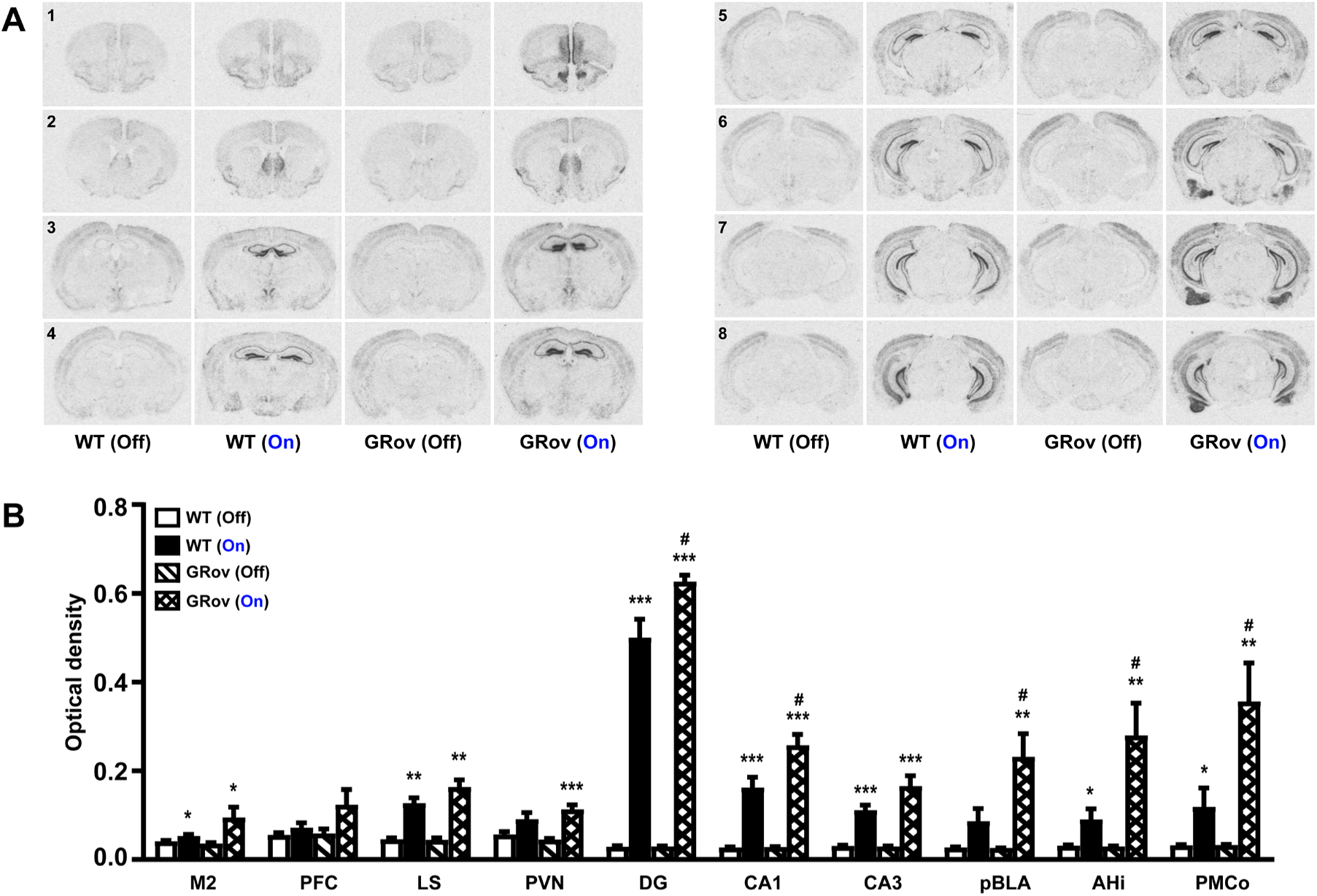
Neuronal activation following optogenetic stimulation in vDG of GRov mice. (*A*) The extent of neuronal activation in the brain following light stimulation in vDG. Relative to dDG, optogenetic activation of vDG led to an increased cFos mRNA expression in more brain regions. Representative images are shown ranging from Bregma 1.98 mm (indicated by the no. 1) through Bregma -3.52 mm (indicated by the no. 8). (*B*) Images were quantitatively analyzed. All listed regions showed increased cFos expression as a result of stimulation in vDG. vDG stimulation resulted in a greater degree of cFos response in DG and CA1 of GRov mice. Activation in vDG resulted in an extensive cFos response in basal posterior amygdaloid complex, including pBLA, AHi, and PMCo. In all three regions neuronal activity was greater in GRov mice than WT. Mean ± SEM. Unpaired two-tailed *t*- test: *P < 0.05, \*\**P* < 0.01, \*\*\**P* < 0.001 versus respective no stimulation group; ^#^*P* < 0.05 versus WT group under the same condition; WT (light off), *n* = 4 mice; WT (light on), *n* = 5 mice; GRov (light off), *n* = 5 mice; GRov (light on), *n* = 5 mice.

Especially remarkable is the differential response of the amygdala in the GRov mice compared to WT. Thus, neuronal activity in pBLA, AHi, and PMCo of GRov mice following vDG stimulation, increased by 7.94, 8.53, and 9.89-fold respectively relative to no stimulation control, (*SI Appendix*, Table S4). In contrast, the neuronal activity in pBLA, AHi and PMCo increased only by 2.23, 1.95, and 2.72-fold respectively in WT animals relative to no stimulation control (*SI Appendix*, Table S4).

Taken together, these results demonstrate that the overexpression of GR in forebrain amplifies the neuronal activity in pBLA, AHi, and PMCo following the stimulation of vDG. They also suggest that the increased and prolonged emotional response to the optogenetic stimulation in vDG of GRov mice engages an extensive neuronal system in the brain, especially in basal posterior amygdaloid complex.

### Glutamatergic/GABAergic Activation Imbalance in vCA1 of GRov Mice Following vDG Activation

We first asked whether GR overexpression alter neuronal activity in a specific population of neurons within the hippocampal formation. Triple fluorescence *in situ* hybridization chain reaction (HCR FISH) labeling was performed on fresh-frozen mouse brain sections to determine the relative number of glutamatergic vs GABAergic cFos-positive neurons in the hippocampus.

In the absence of light stimulation, there was no genotype difference in cFos-based activity in the glutamatergic population within the vDG, vCA1, and vCA3 between GRov and WT mice (*SI Appendix*, Table S5). Following light stimulation in vDG, both GRov and WT mice showed increased cFos activity in the vHC (Fig. 5*A* and *SI Appendix*, Fig. S4), with the most prominent cFos response seen in DG, likely due to the densely packed neurons within the granule cell layer and the fact that this area was the immediate site of illumination by the blue light. Relative to WT, GRov mice showed a comparable number of glutamatergic cFos- positive neurons in vDG and vCA3 (Fig. 5C and *SI Appendix*, Table S5). However, optogenetic stimulation in the vDG caused significantly higher cFos activation of glutamatergic neurons in vCA1 region in GRov relative to WT mice. (Fig. 5 *B*). This was assessed either as the total number of Vglut1^+^/cFos positive neurons (*SI Appendix*, Table S5), or as the ratio of Vglut1^+^/cFos positive neurons relative to the total Vglut1^+^ neuronal population (Fig. 5*C*).

**Fig. 5.**
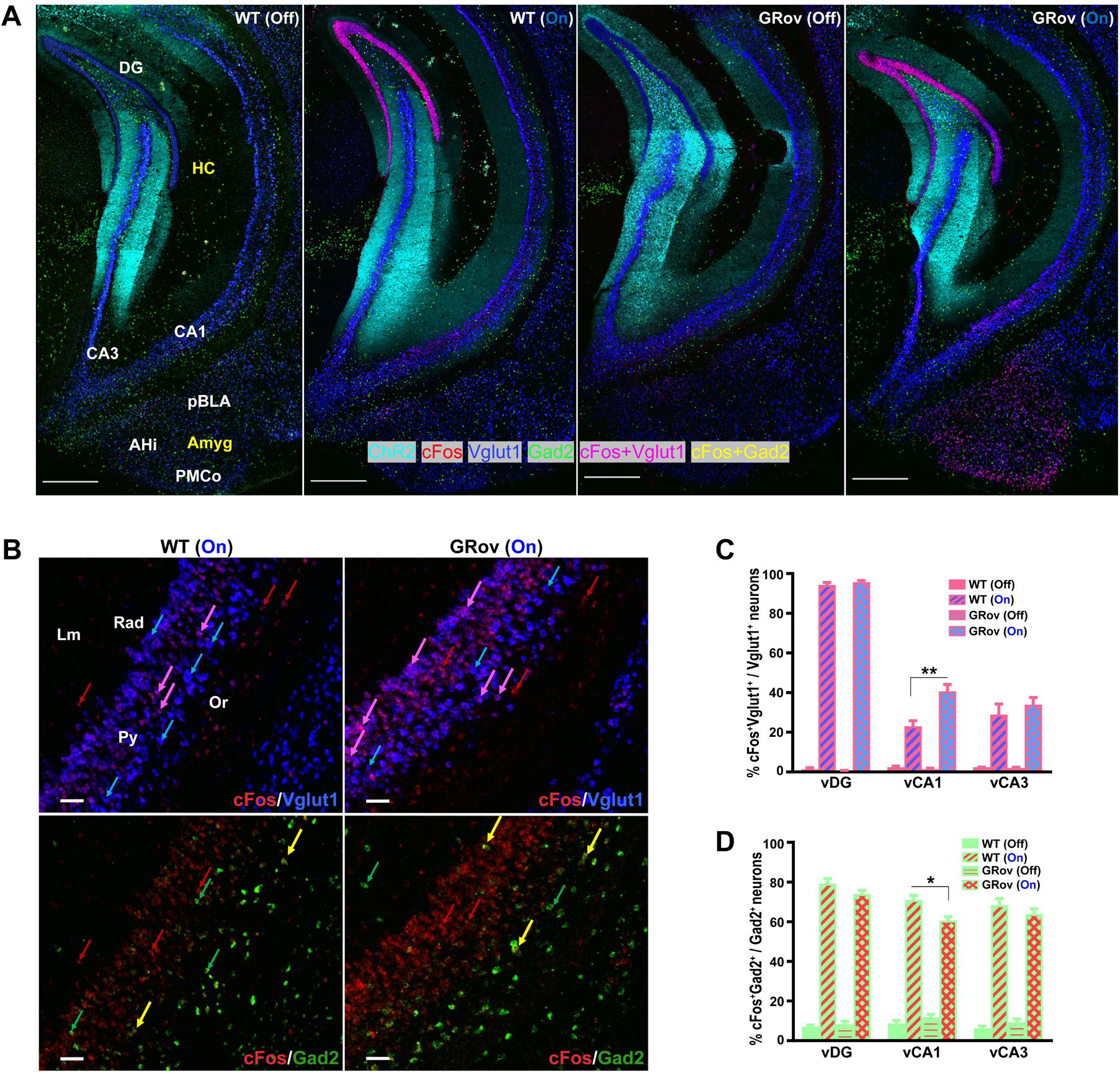
Glutamatergic/GABAergic activation imbalance in vCA1 of GRov mice following optogenetic stimulation in vDG. (*A*) Representative confocal photomicrographs show the increased cFos activity in the ventral hippocampus and the amygdala subregions following blue laser stimulation in vDG of WT and GRov mice compared to no stimulation groups. HCR FISH labeling are represented as cFos (red), Vglut1 (blue), Gad2 (green), ChR2 (cyan), cFos+Vglut1 (magenta), cFos+Gad2 (yellow). Scale bars, 500µm. (*B*) Magnified views of vCA1 show cFos co-labeling with Vglut1^+^ and Gad2^+^. Majority of the cFos+Vglut1 (magenta) labeling can be seen in the pyramidal layer (Py) whereas cFos+Gad2 (yellow) neurons were mostly localized in the stratum oriens (Or), stratum radiatum (Rad) and stratum lacunosum-moleculare (Lm) layers of vCA1. cc- Corpus Callosum. In the representative merged images cFos+Vglut1 and cFos+Gad2 colocalizations are indicated by magenta and yellow arrows, whereas individual labeling of cFos, Vglut1 and Gad2 are indicated by red, blue and green arrows, respectively. Scale bars, 50 µm. (*C*) Histogram shows significant difference in the percent of cFos^+^Vglut1^+^ to total Vglut1^+^ neuronal number density in the vDG, vCA1 and vCA3 following the optogenetic stimulation of vDG in both GRov and WT groups compared to no stimulation groups (not indicated in the graph). There was no genotype difference in cFos-based glutamatergic activity in vDG, vCA1, and vCA3 between GRov and WT mice under no stimulation condition. Mean ± SEM. Unpaired two tailed *t* test: *t*(4) = 0.96, *P* = 0.39 for vDG; *t*(4) = 1.15, *P* = 0.31 for vCA1; *t*(4) = 1, *P* = 0.37 for vCA3; WT (light off), *n* = 3 mice; GRov (light off), *n* = 3 mice. Furthermore, significantly higher cFos-based glutamatergic activity was observed in the vCA1 of stimulated GRov mice compared to stimulated WT animals. Mean ± SEM. Unpaired two tailed *t* test: *t*(7) = 3.83, \*\**P* < 0.01; WT (light on), *n* = 4 mice; GRov (light on), *n* = 5 mice. (*D*) Histogram shows significant difference in the percent of cFos^+^GABA^+^ to total GABA^+^ neuronal number density in the vDG, vCA1, and vCA3 following the optogenetic stimulation of vDG in both WT and GRov mice, compared to no stimulation groups (not indicated in the graph). There was no genotype difference in cFos-based GABAergic activity in vDG, vCA1, and vCA3 between GRov and WT mice under no stimulation condition. Mean ± SEM. Unpaired two tailed *t* test: *t*(4) = 0.72, *P* = 0.51 for vDG; *t*(4) = 1.37, *P* = 0.24 for vCA1; *t*(4) = 1.48, *P* = 0.21 for vCA3; WT (light off), *n* = 3 mice; GRov (light off), *n* = 3 mice. However, in the vCA1 region a significantly lower cFos-based GABAergic activity was observed in the stimulated GRov group compared to stimulated WT group. Mean ± SEM. Unpaired two tailed *t* test: *t*(7) = 3.39, \**P* < 0.05; WT (light on), *n* = 4 mice; GRov (light on), *n* = 5 mice.

A similar analysis was conducted to evaluate the differential activation of GABA neurons in the HC. Basally, there was no genotype difference in vDG, vCA1, and vCA3 subregions between GRov and WT mice (Fig. 5 *B* and *D*). Interestingly, blue light illumination of vDG resulted in a significantly lower cFos response in GABAergic neurons in the vCA1 of GRov mice as compared to WT (Fig. 5 *B* and *D* and *SI Appendix*, Table S5), but not in vDG or vCA3. Thus, the vCA1 of GRov mice has a distinct profile of activation, with a higher proportion of glutamatergic neurons and a lower proportion of GABA neurons exhibiting a cFos response. This suggests that the GRov phenotype may be due to a glutamatergic/GABAergic imbalance in the activity of the vCA1 region.

### Enhanced Glutamatergic Activation in Basal Posterior Amygdaloid Complex of GRov Mice Following vDG Activation

We also characterized the neuronal population activated in the basal posterior amygdaloid complex as a result of vDG stimulation. Both GRov and WT mice showed increased cFos activity in the pBLA, AHi, and PMCo nuclei of amygdala in response to optogenetic stimulation in vDG as compared to no stimulation groups (Fig. 6*A* and *SI Appendix*, Fig. S8 and Table S6). Under basal (no stimulation) conditions, there was no genotype difference in cFos labeling within the glutamatergic population in pBLA, AHi, and PMCo subregions (Fig. 6*D* and *SI Appendix*, Table S6). Following light stimulation in vDG, the majority of activated neurons in pBLA, AHi, and PMCo were identified as glutamatergic in nature (Fig. 6*B* and *SI Appendix*, Table S6). Moreover, GRov mice exhibited significantly higher cFos labeling of glutamatergic neurons in pBLA, AHi, and PMCo subregions. This was evaluated by percentage of activated Vglut1^+^ in total Vglut1^+^ neuronal population (Fig. 6*D*) as well as the total number of activated Vglut1^+^ neurons (*SI Appendix*, Table S6). Thus, 51%, 56%, and 54% of glutamatergic neurons in pBLA, AHi, and PMCo, respectively, were activated in GRov mice as opposed to 13%, 20%, and 23% of glutamatergic neurons in WT (*SI Appendix*, Table S6).

**Fig. 6.**
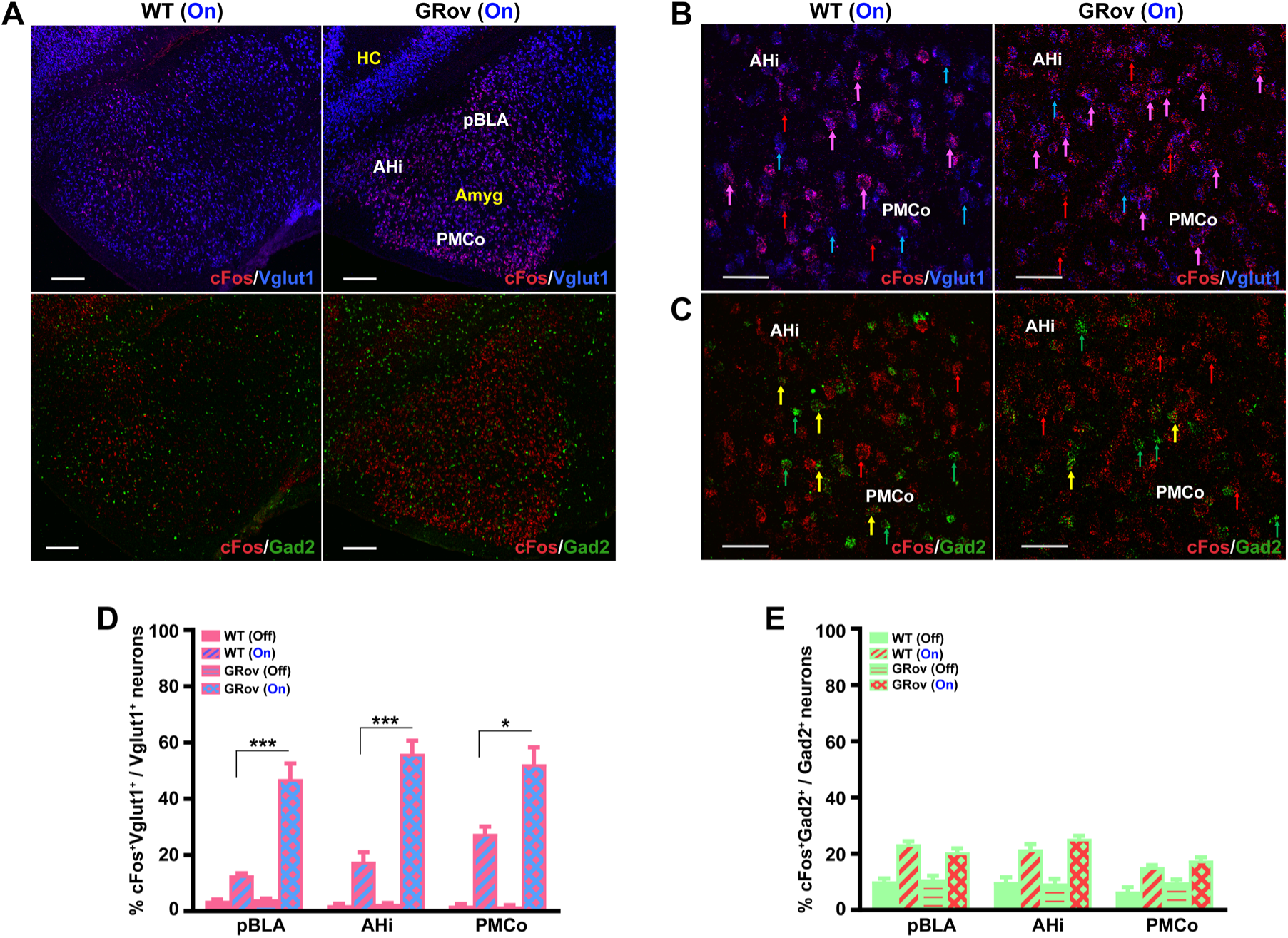
Enhanced glutamatergic activation in basal posterior amygdala of GRov mice following optogenetic activation of vDG. (*A*) Representative confocal images showing the cFos activity in pBLA, AHi, and PMCo following light stimulation in vDG of WT and GRov mice. Greater cFos neuronal density and a predominant colocalization with Vglut1^+^ (magenta) can be seen in GRov-stimulated group compared to WT-stimulated group. cFos (red), Vglut1 (blue), Gad2 (green), cFos+Vglut1 (magenta). Scale bars, 200 µm. (*B*) Magnified views of cFos (red) co-labeling with Vglut1^+^ (blue) show that the majority of activated neurons are Vglut1^+^. Representative cFos+Vglut1 colocalization are indicated by magenta arrows and neurons expressing cFos and Vglut1 are indicated by red and blue arrows, respectively. Scale bars, 50 µm. (*C*) Magnified views of cFos (red) co-labeling with Gad2^+^ (green) show fewer activated (yellow) among the Gad2^+^ population. Representative cFos+Gad2 colocalization are indicated by yellow arrows and neurons expressing cFos and Gad2 are indicated by red and green arrows, respectively. Scale bars, 50 µm. (*D*) Histogram shows significant difference in the percent of cFos^+^Vglut1^+^ to total Vglut1^+^ neuronal number density in the pBLA, AHi, and PMCo following the optogenetic stimulation of vDG in both WT and GRov groups in comparison to the non-stimulated groups (not indicated in the graph). There was no genotype difference in cFos-based glutamatergic activity in pBLA, AHi, and PMCo between GRov and WT mice under no stimulation condition. Mean ± SEM. Unpaired two tailed *t* test: *t*(4) = 0.89, *P* = 0.42 for pBLA; *t*(4) = 0.72, *P* = 0.51 for AHi; *t*(4) = 0.4, *P* = 0.71 for PMCo; WT (light off), *n* = 3 mice; GRov (light off), *n* = 3 mice. Significantly higher cFos-based glutamatergic activity was observed in pBLA, AHi and PMCo of the stimulated GRov group compared to the stimulated WT group. Mean ± SEM. Unpaired two tailed *t* test: *t*(7) = 5.47, \*\*\**P* < 0.001 for pBLA; *t*(7) = 6.39, \*\*\**P* < 0.001 for AHi; *t*(7) = 3.44, \**P* < 0.05 for PMCo; WT (light on), *n* = 4 mice; GRov (light on), *n* = 5 mice. (*E*) Histogram shows significant difference in the percent of cFos^+^GABA^+^ to total GABA^+^ neuronal number density in the pBLA, AHi, and PMCo following the optogenetic stimulation of vDG in both GRov and WT groups compared to no stimulated groups (not indicated in the graph). There was no genotype difference in cFos-based GABAergic activity in pBLA, AHi, and PMCo between GRov and WT mice either basally or under stimulation condition. Mean ± SEM. Unpaired two tailed *t* test. In no stimulation groups, *t*(4) = 0.4, *P* = 0.71 for pBLA; *t*(4) = 0.21, *P* = 0.84 for AHi; *t*(4) = 1.79, *P* = 0.15 for PMCo; WT (light off), *n* = 3 mice; GRov (light off), *n* = 3 mice. In stimulation groups, *t*(7) = 1.64, *P* = 0.14 for pBLA; *t*(7) = 1.83, *P* = 0.11 for AHi; *t*(7) = 1.62, *P* = 0.15 for PMCo; WT (light on), *n* = 4 mice; GRov (light on), *n* = 5 mice.

By contrast, GABAergic neurons in the same amygdala regions did not exhibit any genotype-related differences, either basally or following vDG stimulation (Fig. 6 *C* and *E* and *SI Appendix*, Table S6). These results suggest that overexpression of GR in forebrain selectively amplifies the glutamatergic response within the basal posterior amygdaloid complex in response to activation of the vDG.

### Increased Activation of Calbn1^+^ vCA1 Neurons and Ppp1r1b^+^ pBLA Neurons in GRov Mice Following vDG Activation

Given GRov mice showed a greater and prolonged response to promote risk-taking behavior following the optogenetic activation of vDG, we next investigate whether there are any cell type specific effects within the glutamatergic population both in HC and amygdala, especially in the vCA1 and pBLA subregions which had exhibited the largest differential responses.

The vHC and BLA are implicated in emotion and motivation, playing a critical role in processing both positive and negative emotional events (McKernan et al., 1997; Rogan et al., 1997; Kjelstrup et al., 2002; Rumpel et al., 2005; Tye et al., 2010; Kim et al., 2016; Pi et al., 2020). A known, reciprocal anatomical and functional interaction exists between these regions, and distinct subtypes of glutamatergic neurons have been shown to play distinct functional roles. In particular, the existence of Calbindin1 (Calbn1^+^) in a subset of CA1 hippocampal neurons and of phosphatase 1 regulatory inhibitor subunit 1B-expressing (Ppp1r1b) in a subset of BLA neurons has been implicated in unique functions. We therefore used these two markers to further classify the biochemical identity of those cFos positive glutamatergic neurons via HCR FISH.

In the ventral hippocampal formation, Calbn1^+^ neurons were mainly expressed in the granule cell layer of vDG and pyramidal cell layer of vCA1 (*SI Appendix*, Fig. S9). Distinct and predominant distribution of Calbn1^+^ was observed in the superficial layer of vCA1 relative to deep layer (Fig. 7*A*). Following optogenetic stimulation of vDG, GRov mice displayed increased expression of cFos in Calbn1^+^ neurons in vCA1 compared to WT animals (Fig. 7*A*). This was assessed as the percentage of Calbn1^+^/cFos neurons relative to the total Calbn1^+^ neuronal population (Fig. 7*B*) as well as the total number of Calbn1^+^/cFos neurons (*SI Appendix*, Table S7). There was no difference in Calbn1^+^/cFos neurons in vDG and vCA3 subregions between GRov and WT mice (Fig. 7*B* and *SI Appendix*, Fig. S9 and Table S7). These results demonstrate that the amplified hippocampal response of glutamatergic neurons in GRov mice preferentially engages Calbn1^+^ neurons in the superficial layer of vCA1.

**Fig. 7.**
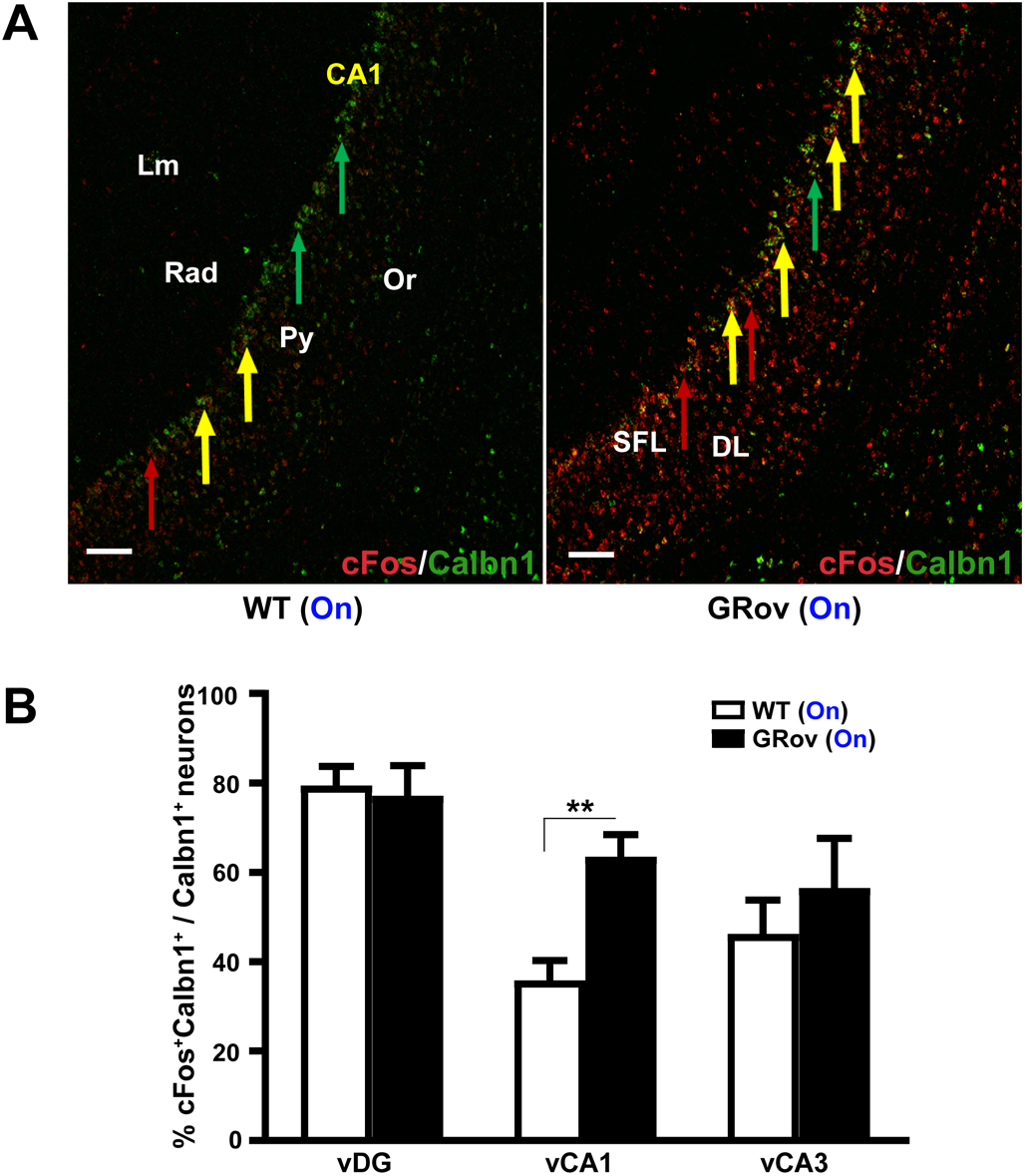
Increased activation of Calbn1-expressing neurons in vCA1 of GRov mice by optogenetic stimulation in vDG. (*A*) Representative confocal photomicrographs showing a greater colocalization (yellow) between Calbn1^+^ neurons (green) and cFos^+^ neurons (red) in the principle (pyramidal) cell layer of vCA1 of GRov- stimulated mice compared to WT-stimulated group following blue laser stimulation in vDG. HCR FISH performed on fresh-frozen brain sections showing distinct and predominant distribution of Calbn1^+^ neurons can be seen in the superficial layer (SFL) of vCA1 compared to deep layer (DL). cFos expression spans across the layers of CA1, including Lm, Rad, Py, Or, whereas Calbn1^+^ expression remains localized mainly within the Py layer. Representative cFos+Calbn1 colocalizations are indicated by yellow arrows, whereas neurons expressing cFos, and Calbn1 are indicated by red and green arrows, respectively. Scale bars, 100µm. (*B*) Histogram shows a significantly higher cFos-based Calbn1 activity in vCA1 of GRov mice following optogenetic stimulation in vDG compared to stimulated WT mice. There was no genotype difference in cFos-based Calbn activity in vDG and vCA3 subregions between GRov and WT mice following optogenetic stimulation in vDG. Mean ± SEM. Unpaired two tailed *t* test: *t*(7) = 0.26, *P* = 0.8 for vDG; *t*(7) = 3.97, \*\**P* < 0.01 for vCA1; *t*(7) = 0.72, *P* = 0.5 for vCA3; WT (light on), *n* = 4 mice; GRov (light on), *n* = 5 mice.

In the basal posterior amygdala, we evaluated the presence or absence of Ppp1r1b, also known as DARPP-32 in the cFos responsive glutamatergic neurons in the pBLA, AHi and PMCo subregions, comparing the GRov to WT mice (Fig. 8*A* and *SI Appendix*, Fig. S10). Here again this was assessed as percentage of Ppp1r1b^+^/cFos neurons relative to the total Ppp1r1b^+^ neuronal population (Fig. 8*B*) as well as the number of activated Ppp1r1b^+^/cFos neurons (*SI Appendix*, Table S8). Remarkably, the number Ppp1r1b^+^/cFos neurons in pBLA subregion is 4.31- fold higher in GRov mice when compared with WT mice (*SI Appendix*, Table S8). These results demonstrate that the amplified response of glutamatergic neurons in GRov mice preferentially engages Ppp1r1b^+^ neurons in the basal posterior amygdala subregions, especially in pBLA.

**Fig. 8.**
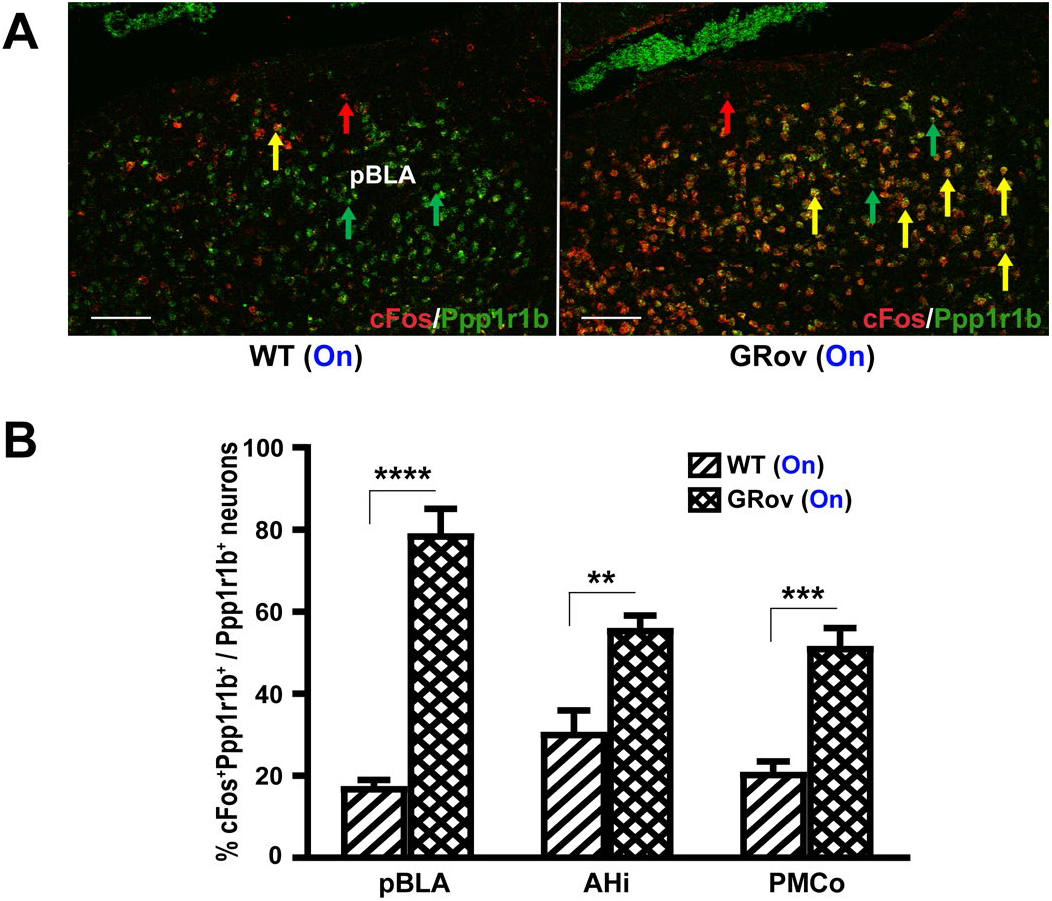
Increased activation of Ppp1r1b-expressing neurons in pBLA of GRov mice by optogenetic activation in vDG. (*A*) Representative confocal photomicrographs of pBLA showing a greater colocalization (yellow) of Ppp1r1b^+^ neurons (green) with cFos^+^ neurons (red) in the GRov-stimulated mice compared to WT-stimulated group following blue laser stimulation in vDG. HCR FISH performed on fresh-frozen brain sections showing representative cFos+Ppp1r1b colocalizations (yellow arrows) and neurons expressing cFos and Ppp1r1b with red and green arrows, respectively. Scale bars, 100µm. (*B*) Histogram shows a significantly higher cFos-based Ppp1r1b^+^ activity in pBLA, AHi, and PMCo of GRov mice following optogenetic stimulation in vDG compared to stimulated WT mice. Mean ± SEM. Unpaired two tailed *t* test: *t*(7) = 8.93, \*\*\*\**P* < 0.0001 for pBLA; *t*(7) = 4.34, \*\**P* < 0.01 for AHi; *t*(7) = 5.66, \*\*\**P* < 0.001 for PMCo; WT (light on), *n* = 4 mice; GRov (light on), *n* = 5 mice.

## Discussion

In this study, we sought to identify the neural circuitry associated with increased emotional reactivity. To this end, we used our well characterized animal model of emotional lability- a mouse that overexpresses GR only in the forebrain and especially in the hippocampus. This animal has a normal basal neuroendocrine stress profile but exhibits rapid swings in emotional responses to various contexts and psychoactive drugs. In this study, we demonstrated similar emotional lability with optogenetic stimulation and uncovered a specific neural circuit associated with this amplified emotional response. Our findings are: a) Optogenetic stimulation of the DG leads to greater risk taking behavior (percent entries into the open arms and percent time in the open arms of the EPM) in both WT and GRov mice; b) the impact of the optogenetic stimulation is amplified in the GRov mice and this is especially evident with vDG stimulation which led to both greater amplitude and longer duration of the risk taking behavior in the GRov animals; c) stimulation of both dDG and vDG elicited distinct neural activation across both groups of mice as measured by cFos, but effects of vDG stimulation were more widespread; d) the enhanced GRov behavioral response is coupled with intense neuronal activation in hippocampal CA1 region and in the basal posterior amygdala- notably pBLA, AHi and PMCo; e) the enhanced neural response in GRov mice was selective to glutamatergic neurons. This was seen in the CA1 as well as in the pBLA, AHi, and PMCo regions of the amygdalar complex. In addition, there was decreased activation of GABAergic neurons in the CA1 of GRov relative to WT; f) only specific subpopulations of glutamatergic neurons were differentially activated by GR overexpression- Calbn1^+^ vCA1 neurons and Ppp1r1b^+^ pBLA neurons. Thus, a molecularly distinct component of the hippocampal-amygdala circuit is modified by early life changes in the stress system and is associated with a propensity for greater emotional lability.

In the present study, GRov mice showed greater risk-taking behavior following optogenetic activation of either dDG or vDG when compared to WT, with the effect being greater and more prolonged following vDG activation. The increased exploratory behavior in the EPM following photostimulation of the DG is consistent with a previous finding that elevating the activity of granule cells in the dorsal DG led to a dramatic increase in exploration (Kheirbek et al., 2013). It should be noted that the enhanced emotional reactivity in GRov mice following optogenetic stimulation in dDG or vDG is independent of their increased exploration, as GRov and WT mice showed comparable exploratory activity during the photostimulation epoch in the EPM test. Taken together, these behavioral results indicate that, in response to activation of vDG, overexpression of GR in forebrain promotes greater emotional swings, modifying both the magnitude and the duration of responsiveness to the stimulus.

We then investigated the potential neuronal mechanisms associated with these behavioral profiles by using cFos mRNA as a cellular activity marker. In both GRov and WT mice, photostimulation of dDG or vDG activated numerous key brain regions which have been implicated in the modulation of emotional reactivity. These include cerebral cortex (PFC, prelimbic, infralimbic and cingulate), M1/2 and S1 cortex, LS, BNST, thalamus, PVN, hypothalamic nuclei, HC, and amygdaloid complex.

Optogenetic stimulation of dDG specifically induced higher neuronal activity in LS, DG, CA1, and pBLA of GRov mice compared to WT animals. The extensive connection between septum and HC forms the septohippocampal pathway. The septum contains cholinergic, GABAergic, and glutamatergic neurons that project to the HC via the fimbria-fornix (Freund and Antal, 1988; Kiss et al, 1990). The HC projects to the septum mainly via glutamatergic afferents (Niewiadomska et al., 2009). A large body of evidence indicates that the septohippocampal pathway plays a role in regulating anxiety-related behaviors. For instance, chemogenetic activation of LS-projecting ventral hippocampus neurons decreased anxiety whereas inhibition produced anxiety-promoting effects (Parfitt et al., 2017).

In contrast, vDG activation in GRov mice led to robust neuronal activation in DG, CA1, and basal posterior amygdala subregions (pBLA, AHi, and PMCo) as evidenced by the intense cFos radioactive-ISH signal. Moreover, multiplexed-HCR FISH studies revealed activation of a significant population of glutamatergic neurons in vCA1, pBLA, AHi, and PMCo of GRov mice. Together these findings indicate that GR overexpression greatly amplifies the responsiveness of a neural circuit that comprises the vCA1 and the basal posterior amygdala to vDG activation.

What are the likely elements of this neural circuit? There is ample evidence for the existence of a direct reciprocal anatomical connection between the amygdala and the hippocampus. Extensive anterograde and retrograde tracing data indicates that the amygdaloid complex is reciprocally connected to the hippocampus and the surrounding cortex. Projections from the hippocampal formation to the amygdala arise solely from CA1 (Pitkänen et al., 2006; Kishi et al., 2006; McDonald et al., 2017), which receive outputs from the tri-synaptic circuit. Electron microscopic studies have shown that the projections from the vCA1 form asymmetrical (excitatory) synapses with the distal dendrites and spines of the pyramidal cells in the basolateral complex (Müller et al., 2012). The projections from the nuclei of amygdala largely reciprocate the hippocampal projections to the amygdaloid complex. The interaction between basal amygdala and ventral hippocampus has been associated with emotion and motivation, and processing of both positive and negative emotion-associated events (McKernan et al., 1997; Rogan et al., 1997; Kjelstrup et al., 2002; Rumpel et al., 2005; Tye et al., 2010). Here, we found that GRov mice display enhanced risk-taking behavior in conjunction with the amplified activation of this neural circuit.

Using HCR FISH, we have shown that the majority of cFos-based activated neurons in the pyramidal cell layer of vCA1 (PCL-vCA1) and basolateral amygdala are glutamatergic. Moreover, the increased activation in GRov mice is almost exclusively glutamatergic. Indeed, in the context of the CA1, we observed a decrease in GABA neurons being activated in GRov implying a strong bias towards glutamatergic activity within the CA1-BLA circuit.

We then investigated the cell-type specificity within these predominant patterns of cFos^+^ neurons in PCL-vCA1 and pBLA subregions, which might help explain the increased risk-taking behavior following the optogenetic activation of vDG in GRov mice. GRov mice exhibited increased activation of Calbn1^+^ neurons in vCA1 and increased activation of Ppp1r1b^+^ neurons in pBLA following optogenetic activation of vDG. The basolateral amygdala consists of two types of organized excitatory pyramidal neurons, magnocellular and parvocellular, segregated respectively into the anterior and posterior subregions (Pitkänen et al., 1997; McDonald, 1982). Manipulating the neuronal activity in anterior BLA and pBLA can elicit negative and positive emotional behaviors, respectively (Kim et al., 2016). Ppp1r1b^+^ neurons located in pBLA respond to positive valence stimuli and control appetitive behaviors and memories (Kim et al., 2016). A recent study has shown that fear extinction memory engrams are formed and stored within the basolateral amygdala Ppp1r1b^+^ neurons and that these engram cells are necessary for suppressing the original fear memory (Zhang et al., 2020). This then points to Ppp1r1b^+^ neurons as favoring approach behaviors.

Additional work implicates the pBLA-CA1 in the selection of approach vs avoidance behaviors. Excitatory monosynaptic upscaling between pBLA and vCA1 is associated with “learned hopefulness”, and optogenetic disruption of pBLA– vCA1 inputs abolishes the effects of “learned hopefulness” and impairs synaptic plasticity (Yang et al., 2016). Moreover, photostimulation of the pBLA–vCA1 excitatory neurons has an anxiolytic effect in mice, promoting approach behaviors during conflict exploratory tasks (Pi et al., 2020). This stands in contrast to the effects of activating the anterior BLA-vHPC inputs which increases avoidance behavior and exerts anxiogenic effects (Felix-Ortiz et al., 2013), Given our findings, the role of calbindin 1 positive neurons within the CA1 is of particular interest. Recent work has focused on the conflict stages of the EPM task and showed that photostimulation of vCA1 Calbn1^+^ excitatory neurons at the decision-making zones promotes approach with fewer retreats (Pi et al., 2020). It is therefore reasonable to suggest that optogenetic activation of the vDG by selectively increasing the activity of Calbn1^+^ neurons in vCA1 and Ppp1r1b^+^ neurons in pBLA resulted in greater exploratory behavior in both the WT and GRov mice. It should be recalled that our GRov mice have more baseline anxiety. However, photostimulation of their vDG led to greater activation of the Calbn1^+^ and Ppp1r1b^+^ neurons in the vCA1-BLA circuit. This combination likely mediates, at least in part, the broad swing in emotionality observed in the EPM for the GRov mice.

While exploratory anxiety strongly engages the ventral hippocampus, other emotional behaviors that also exhibit greater swings in the GRov mouse, such as the antidepressant response in the forced swim test, or the response to psychostimulants, likely engage other elements of affective circuitry. It would be of great interest to ascertain whether the vCA1-BLA circuit also contributes to these manifestations of greater lability.

Finally, it should be recalled that the GRov phenotype is established early in development and affects responsiveness to a broad range of emotional behaviors throughout the animal’s life. High density of endogenous GR has been shown to render animals highly susceptible to alterations in glucocorticoid levels— i.e., highly sensitive to stress (Jacobson and Sapolsky, 1991). Overexpression of GRov mimics a condition of high stress exposure during a key organizational stage of the brain. This is especially relevant to the dentate gyrus which continues to develop and show great plasticity postnatally and is profoundly affected by glucocorticoid manipulations (Gould et al, 1991; McEwen et al, 2016). We therefore propose that perturbations of the stress system in general, and glucocorticoid receptors in particular, during a critical developmental period retunes affective circuitry to render individuals more emotionally reactive, including altering the responsiveness of the vCA1-BLA circuit characterized here. It would be of great interest to ascertain whether this circuitry is implicated in human conditions, such as bipolar disorder, that involve dysregulation of emotionality and greater propensity for mood swings.

## Materials and Methods

### Animals

The generation of transgenic mice overexpressing GR specifically in forebrain was described previously (Wei et al., 2004). GRov mice were established by breeding founders and their progeny to C57BL/6J mice. Mice were housed in a climate-controlled environment at 22°C with a 14:10 light/dark schedule of lights on between 5:00 and 19:00 and *ad libitum* access to standard lab pellet food (5L0D-PicoLab, Brentwood, MO) and water. All transgenic mice are maintained as hemizygotes. All experiments were performed with male GRov mice and their WT littermates. Mice were between 3 and 6 months old at the time of testing. All experiments were conducted in accordance with the principles and procedures outlined in the National Institutes of Health guidelines for the Care and Use of Animals and were approved by the IACUC at University of Michigan.

### Stereotactic Surgery and Virus Vector Injection

Mice were anesthetized with isoflurane and placed into a stereotaxic apparatus (David Kopf, CA). After exposing the skull via a small incision, a small hole was drilled. 0.8 µl of AAV5- CaMKIIa-ChR2(H134R)-eYFP virus (UNC vector core, Chapel hill, NC) was injected unilaterally in the right dDG (coordinates, Bregma: AP: -1.7 mm, DV: 2.05 mm, ML: 0.63 mm) or vDG (coordinates, Bregma: AP: -3.3 mm, DV: 2.25 mm, ML: 2.1 mm) using a microsyringe pump injector (model UMP3, World precision instruments, CA) that controlled a NanoFil syringe (World precision instruments) to generate an injection rate of 100 nl per min. Following injection, the syringe was slowly withdrawn after waiting 10 min. Immediately after withdrawal of the injection syringe, an optical fiber (200 µm core, NA 0.37, Doric lenses, Canada) was implanted over the dDG (AP: -1.7 mm, DV: 1.75 mm, ML: 0.63 mm) or vDG (AP: -3.3 mm, DV: 1.95 mm, ML: 2.1 mm). Mice were singly housed after surgery and allowed to recover for 3 weeks before experiments.

### Elevated Plus Maze Test

The elevated plus maze (EPM) consisted of four arms (27 × 6 cm) arranged in a plus form, and elevated 50 cm above the floor. Two opposing arms were surrounded with 15-cm-high clear Plexiglas walls (closed arms) while the other arms were devoid of walls (open arms). The light intensity measured in the open arms was 180 Lux. Tests were conducted at least 3-week post-surgery during the same circadian period (7:00 – 12:00). Mice were handled and attached to the optical fiber without photostimulation for 3 days before behavioral test. On testing day, mice were connected to the optical fiber and placed in an empty mouse cage for 2 min, then mice were placed in the center of the EPM, facing an open arm to begin the test. Sessions lasted for 18 min consisting of three 6 min epochs: blue laser off (Off1), blue laser on (On), and blue laser off (Off2). During the blue laser (473 nm, Vortran Laser Technology, CA) stimulation epoch, the laser parameters were 10 ms light pulses at 10 Hz for 30 s every minute for 6 min with 2.7-3.1 mW light power on the tip of fiber. An entry was defined as having 4 paws in an arm of the plus maze.

### cFos mRNA Expression Following Optogenetic Stimulation

For cFos mRNA response experiments, mice were attached to the optical fiber without photostimulation and habituated to an open field chamber (36 X 36 cm) for 3 consecutive days before laser stimulation. Mice received 3 min of either blue laser stimulation (10 ms pulses at 10 Hz for 30 s every minute for 3 min with 2.7-3.1 mW on the tip of fiber) or no laser stimulation in the same open field chamber, then returned to their home cage. Mice were killed by rapid decapitation 30 min after the onset of laser stimulation and their brains were removed, snap-frozen in 2- methylbutane, and stored at −80°C until sectioned.

*In situ* hybridization (ISH) methodology was performed as previously described (Wei et al., 2018). The cFos probe was a 667-bp fragment directed against the mouse cFos mRNA. cFos cDNA segment was extracted, subcloned in Bluescript SK (Stratagene), and confirmed by nucleotide sequencing. The probe was labelled in a reaction mixture of 1 μg of linearized plasmid, 1× transcription buffer (Epicentre Technologies), 125 μCi of ^35^S-labeled UTP, 125 μCi of ^35^S-labeled CTP, 150 μM ATP and GTP, 12.5 mM DTT, 1 μL RNase inhibitor, and 1.5 μL RNA polymerase. After hybridizing the labelled probe to the sectioned tissue, sections were thoroughly washed and exposed to Kodak BioMax MR Scientific Imaging Film (Sigma Aldrich). Radioactive signals were quantified using computer-assisted optical densitometry software ImageJ (National Institutes of Health).

### Fluorescence In Situ Hybridization Chain Reaction (HCR FISH)

Split-initiator DNA probes (*version* 2.0) were designed in our lab (Choi et al., 2018; Kumar et al., 2021) (*SI Appendix*, Table S9) and synthesized by Integrated DNA Technologies (Coralville, Iowa, USA). DNA hairpins conjugated with AlexaFluor-546 (AF-546), AF-594 and AF-647 were purchased from Molecular Instruments, Inc. (Los Angeles, California, USA). Fresh-frozen brains were cryosectioned to get 12 sections (20 µm thickness) between ‘Bregma -2.92 mm to -3.40 mm’ for vHC and between ‘Bregma -2.18 mm to -3.40 mm’ for amygdala and used for HCR FISH. We optimized the HCR FISH method as described previously (Choi et al., 2014, 2018; Kumar et al. 2021). Briefly, sections were immersion fixed in 4% paraformaldehyde-saline, washed with 5x sodium chloride/sodium citrate/0.01% tween-20 (SSCTw) buffer for 3 times, 5 min each, then acetylated in 0.1M triethanolamine, pH 8.0 with 0.25% vol/vol acetic anhydride solution for 10 min. After rinsing with ddH2O, sections were de-lipidated in -20^◦^C chilled acetone: methanol (1:1) for 5 min, washed with 5xSSCTw, and equilibrated in hybridization buffer (30% deionized formamide, 5x SSC, 9 mM citric acid (pH 6.0), 0.5 mg/ml yeast tRNA, 1x Denhardt’s solution, 10% dextran sulfate, 0.1% Tween 20) for 60 min, then incubated in hybridization buffer containing 10nM initiator-labeled probes at 37°C for 16 hrs. Following hybridization, sections were washed at 37°C with probe wash buffer (30% formamide, 5x SSC, 0.1% tween 20) for 3 times and twice with 5x SSCTw for 15 min each. Sections were then equilibrated in amplification buffer for 60 min (5x SSC, 10% dextran sulfate, 0.1% Tween 20). Fluorophore-labeled hairpins were diluted separately from 3 µM stock to a 2.25 µM final concentration in 20x SSC, heated at 90°C for 90 seconds, and then snap-cooled to room temperature for 30 min in the dark. Snap-cooled hairpins were further diluted to 60nM final concentration in amplification buffer. Sections were incubated in amplification buffer with hairpins for 16 hrs at room temperature. Finally, tissues were washed in 5x SSCTw twice for 30 min and mounted with Vectashield antifade mounting medium (Vector Laboratories, Inc., Burlingame, CA 94010, United States).

### Confocal Microscopy

Image stacks were acquired using an Olympus Fluoview- 3000 confocal microscope and consisted of 5 channels, DAPI+eYFP+cFos+Vglut1+Gad2 or DAPI+eYFP+cFos+Calbn1+Ppp1r1b. For quantitative colocalization analysis, a 10x magnification objective lens (Olympus UPLSAPO10X2, numerical aperture (N.A.) 0.4 / working distance (W.D.) 2.2 mm) was used to acquire image-stacks (xy-dimension 1.59 µm/pixel x 1.59 µm/pixel and z-step of 4.5 µm; 4-6 z-slices per stack). High magnification representative images were acquired using 20x (air, Olympus UCPLFLN20X, N.A.:0.7/W.D.:0.80- 1.80mm) or 40x (silicone oil immersion, Olympus UPLSAPO40XS, N.A.- 1.25/W.D.- 0.3mm) objectives. Image acquisition settings mainly the PMT Voltage and laser transmissivity were selected for optimal pixel saturation to avoid excessive or weak signal and kept constant across the study.

### Image Processing and Analysis

Images from 3-4 sections per animal consisting of whole ROIs, HC and amygdala (*n* = 3 for controls, *n* = 4-5 for stimulated groups) were processed and quantified. Open-source ImageJ/Fiji software (Schneider et al., 2012; Schindelin et al., 2012) was used for visualization, processing and quantitation. Region of interests (ROIs), CA1, CA3, DG, pBLA, AHi and PMCo were drawn using the free-hand tool and managed in the ROI manager for quantitation of neuronal number from each channel. Two atlases, ‘the mouse brain in stereotaxic coordinates’ by Paxinos & Franklin (Academic Press, 2001), and ‘the Allen Institute mouse brain atlas’ were followed for ROI drawings. Mean data of sections from each animal was then used for statistical analysis. Images were processed globally as 3D stacks, first filtered using an anisotropic diffusion method which effectively preserves strong edges and reduces the background, followed by Gaussian blur to concentrate the signal towards the center of each cell. DAPI signal (nuclear marker) was selectively processed for maxima finder and spot segmentation to create properly segmented selections of individual cells. These selections were used as masks on the filtered grey channel images and segmented with Otsu-thresholding algorithm. After segmentation of two images (e.g., to analyze colocalization between cFos and Vglut1), the image calculator tool was used to generate the intersection (AND operator) image containing common pixel data. Analyze particles tool (appropriate size and circularity filter set for each signal type) was used to count objects from the AND generated image (e.g., cFos^+^Vglut^+^) and the two individual channels (e.g., cFos and Vglut1) for each ROI (e.g., CA1, CA3, pBLA etc.). Images were processed similarly for Gad2, Calbn1 and Ppp1r1b channels. Using maximum thresholding values, area of each ROI was also calculated and then used to estimate the neuronal number per unit area (number density). Percent of colocalized cells were then calculated and used for histograms. Statistical analysis was performed on both percent and raw number density data.

### Statistical Analysis

Statistical analyses were performed using Prism 9.2.0 (GraphPad) software. *P* < 0.05 was considered statistically significant.

## ACKNOWLEDGMENTS

This work was supported by the Hope for Depression Research Foundation, the Pritzker Neuropsychiatric Research Consortium, the Office of Naval Research (Grant # ONR-00014-19-1-2149. We thank Dr. Karl Deisseroth (Stanford University, Department of Bioengineering) for the gift of AAV5-CaMKIIa-ChR2(H134R)-eYFP virus and AAV5-CaMKIIa-mCherry virus. We thank Dr. Cortney Turner for helpful discussion of the manuscript, Limei Zhang and Mathew Foltz for technical assistance.

## COMPETING FINANCIAL INTERESTS

The authors declare no conflict of interest.

## Notes

### Competing Interest Statement

The authors have declared no competing interest.

