## supplementary file for "High emotional reactivity is associated with activation of a molecularly distinct hippocampal-amygdala circuit modulated by the glucocorticoid receptor"

* Equal First Authors
** Equal Last Authors

[1]To whom correspondence should be addressed:

Huda Akil, PhD
Michigan Neuroscience Institute
University of Michigan
205 Zina Pitcher Place, Ann Arbor, MI 48109


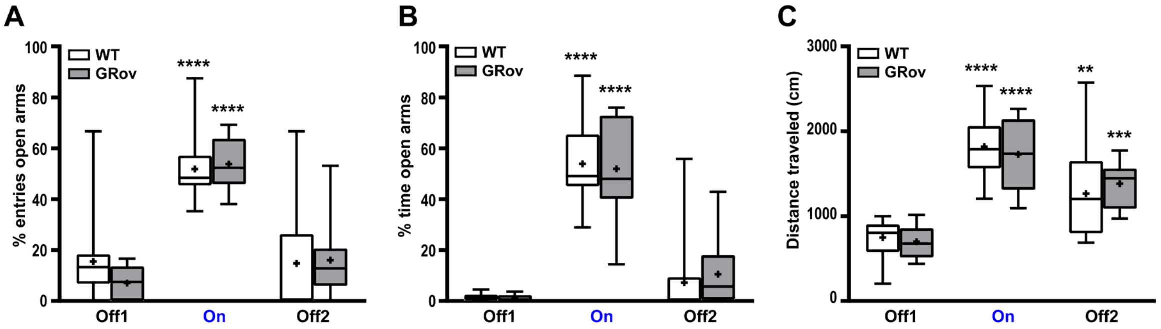


**Fig. S1.** Optogenetic activation of dDG induced robust and reversible increase in risk-taking behavior in the EPM test in both GRov and WT mice. (*A*) Repeated measures two-way ANOVA revealed there was no genotype × light interaction, $F(2, 42) = 0.9$, $P = 0.41$, for entries into the open arms in EPM test. Repeated-measures one-way ANOVA revealed that blue light stimulation in dDG led to increased entries into the open arms in GRov [$F(2, 18) = 67.61$, $P < 0.0001$; ****$P < 0.0001$ versus pre-stimulation] and WT [$F(2, 24) = 19.09$, $P < 0.0001$; ****$P < 0.0001$ versus pre-stimulation]. (*B*) Repeated measures two-way ANOVA revealed there was no genotype × light interaction, $F(2, 42) = 0.27$, $P = 0.77$, for time spent in open arms in EPM. Repeated-measures one-way ANOVA revealed that blue light stimulation in dDG led to increased time spent in the open arms in GRov [$F(2, 18) = 38.5$, $P < 0.0001$; ****$P < 0.0001$ versus pre-stimulation] and WT [$F(2, 24) = 82.78$, $P < 0.0001$; ****$P < 0.0001$ versus pre-stimulation]. (*C*) Repeated measures two-way ANOVA showed no genotype × light interaction, $F(2, 42) = 0.52$, $P = 0.6$, for total distance traveled in EPM. Repeated-measures one-way ANOVA revealed that blue light stimulation in dDG led to increased distance traveled in EPM in GRov [$F(2, 18) = 27.69$, $P < 0.0001$; ***$P < 0.001$, ****$P < 0.0001$ versus pre-stimulation] and WT [$F(2, 24) = 23.51$, $P < 0.0001$; **$P < 0.01$, ****$P < 0.0001$ versus pre-stimulation] and this increase in general exploration extended after the termination of blue light stimulation. Box plots show median, mean (+), lower and upper quartiles (boxes), and minima and maxima (whiskers). GRov, $n = 10$ mice; WT, $n = 13$ mice.

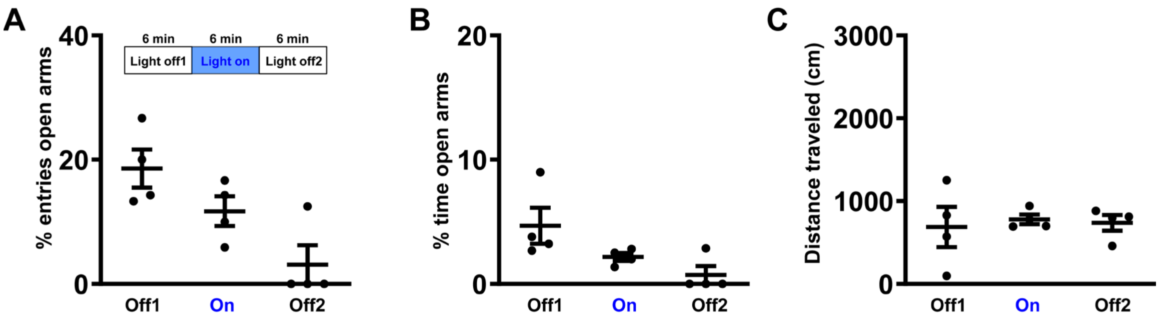


**Fig. S2.** Normal habituation behavior following control virus injection in dDG. (*A* and *B*) Blue light illumination of the dDG in mice with control virus microinjection in dDG didn't trigger any increases in entries into the open arms or time spent in the open arms of the EPM. Repeated-measures one-way ANOVA revealed that these mice displayed a normal habituation behavioral phenotype across testing epochs. $F(2, 6) = 12.92$, $P = 0.007$ for entries into open arms; $F(2, 6) = 3.49$, $P = 0.1$ for time spent in open arms; $n = 4$ mice. (C) Mice with control virus injection in dDG displayed comparable total distance travelled in EPM across testing epochs. $F(2, 6) = 0.08$, $P = 0.92$; $n = 4$ mice.

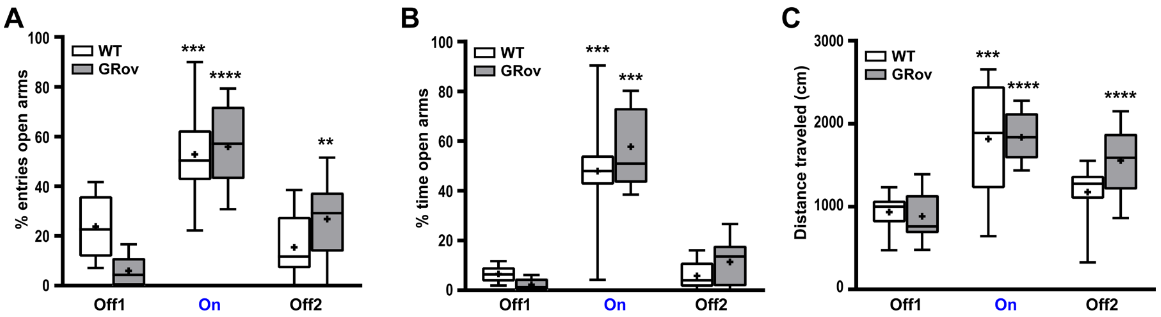


**Fig. S3.** Optogenetic activation of vDG induced robust increase in risk-taking behavior in the EPM test in both GRov and WT mice. (*A*) Repeated measures two-way ANOVA revealed a significant genotype × light interaction, $F(2, 34) = 6.45$, $P < 0.01$, for entries into the open arms in EPM test. Repeated-measures one-way ANOVA revealed that blue light stimulation in vDG led to increased entries into the open arms in GRov [$F(2, 20) = 36.92$, $P < 0.0001$; **$P < 0.01$, ****$P < 0.0001$ versus pre-stimulation] and WT [$F(2, 14) = 14.82$, $P < 0.001$; ***$P < 0.001$ versus pre-stimulation]. (*B*) Repeated measures two-way ANOVA revealed there was no genotype × light interaction, $F(2, 34) = 1.77$, $P = 0.19$, for time spent in open arms in EPM test. Repeated-measures one-way ANOVA revealed that blue light stimulation in vDG led to increased time spent in the open arms in GRov [$F(2, 20) = 76.5$, $P < 0.0001$; ****$P < 0.0001$ versus pre-stimulation] and WT [$F(2, 14) = 30.28$, $P < 0.0001$; ****$P < 0.0001$ versus pre-stimulation]. (*C*) Repeated measures two-way ANOVA revealed there was no genotype × light interaction, $F(2, 34) = 2.61$, $P = 0.09$, for total distance traveled in EPM. Repeated-measures one-way ANOVA revealed that blue light stimulation in vDG led to increased distance traveled in EPM in GRov [$F(2, 20) = 31.26$, $P < 0.0001$; ****$P < 0.0001$ versus pre-stimulation] and WT [$F(2, 14) = 14.82$, $P < 0.001$; ***$P < 0.001$ versus pre-stimulation]. Box plots show median, mean (+), lower and upper quartiles (boxes), and minima and maxima (whiskers). GRov, $n = 11$ mice; WT, $n = 8$ mice.

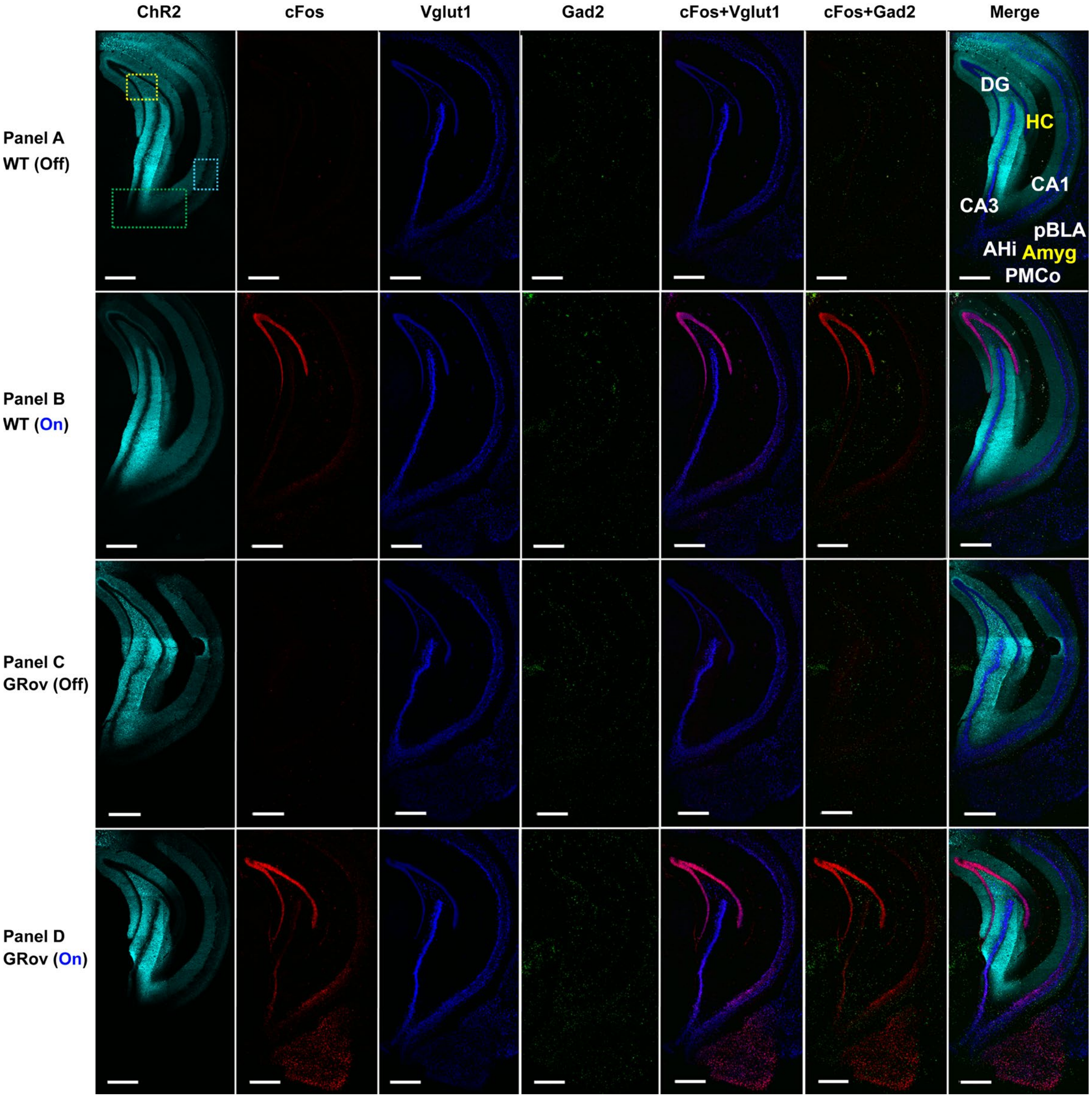


**Fig. S4.** Representative confocal images showing the increased cFos activity in the ventral hippocampus following light stimulation in vDG of WT and GRov mice (panels B and D) compared to no stimulation groups (panels A and C). Split channels of the representative images are shown as ChR2 (cyan), cFos (red), Vglut1 (blue), Gad2 (green) and merged color combinations cFos+Vglut1 (magenta), cFos+Gad2 (yellow) and merged (all channels) of the representative images are shown. Higher cFos activity co-localized with Vglut1$^{+}$ glutamatergic and Gad2$^{+}$ GABAergic neurons in the hippocampus and amygdala subregions of both WT and GRov animals (panels B, D) following the optogenetic stimulation in vDG. Scale bars, 500 µm. Insets-refer to *SI Appendix*, figures S5, S6, and S7 showing zoomed-views of the co-labeling in vDG, vCA3-vCA1, and vCA1 respectively.

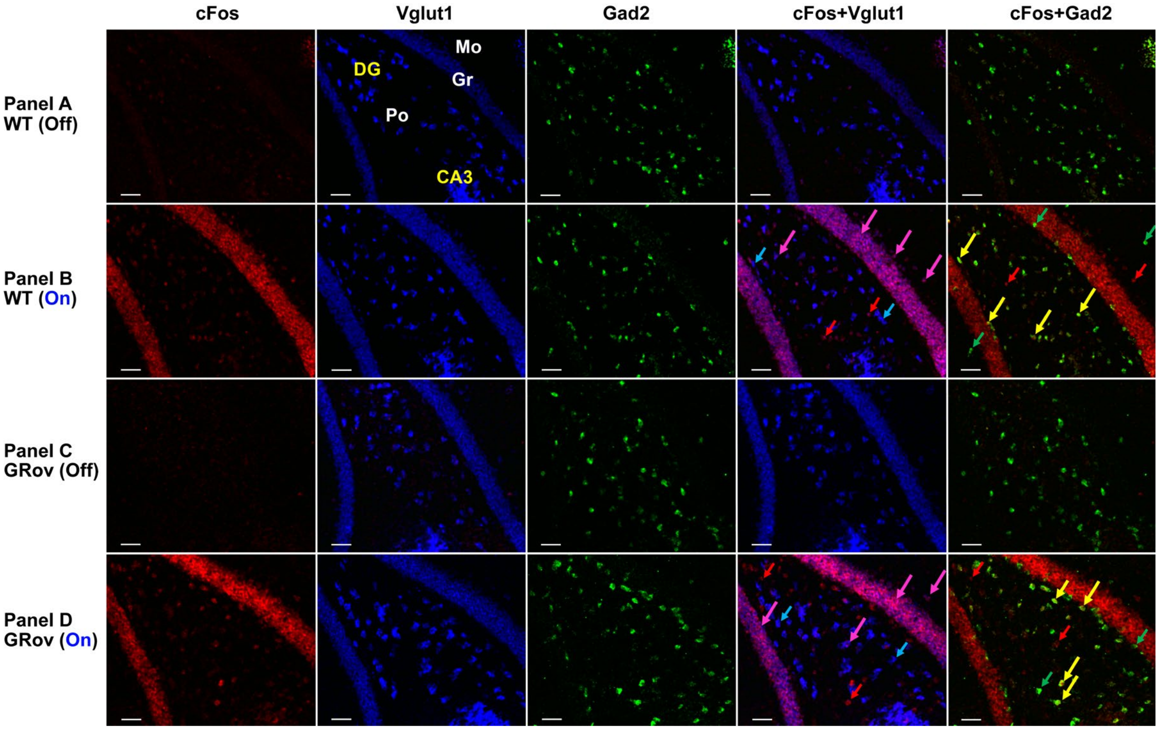


**Fig. S5.** Magnified views of DG of ventral hippocampus showing cFos co-labeling with Vglut1$^{+}$ and Gad2$^{+}$ in the regions corresponding to the insets shown in Fig. S4. Most of the cFos+Vglut1 (Magenta) labeling can be seen in the granule layer (Gr) whereas cFos+Gad2 (Yellow) neurons were mainly localized in the polymorph layer (Po). Representative merged-color columns show cFos+Vglut1 colocalization (magenta arrows) and cFos+Gad2 colocalization (yellow arrows) and cFos, Vglut1 and Gad2 expressing neurons as red, blue, and green arrows, respectively. Scale bars, 50 μm.

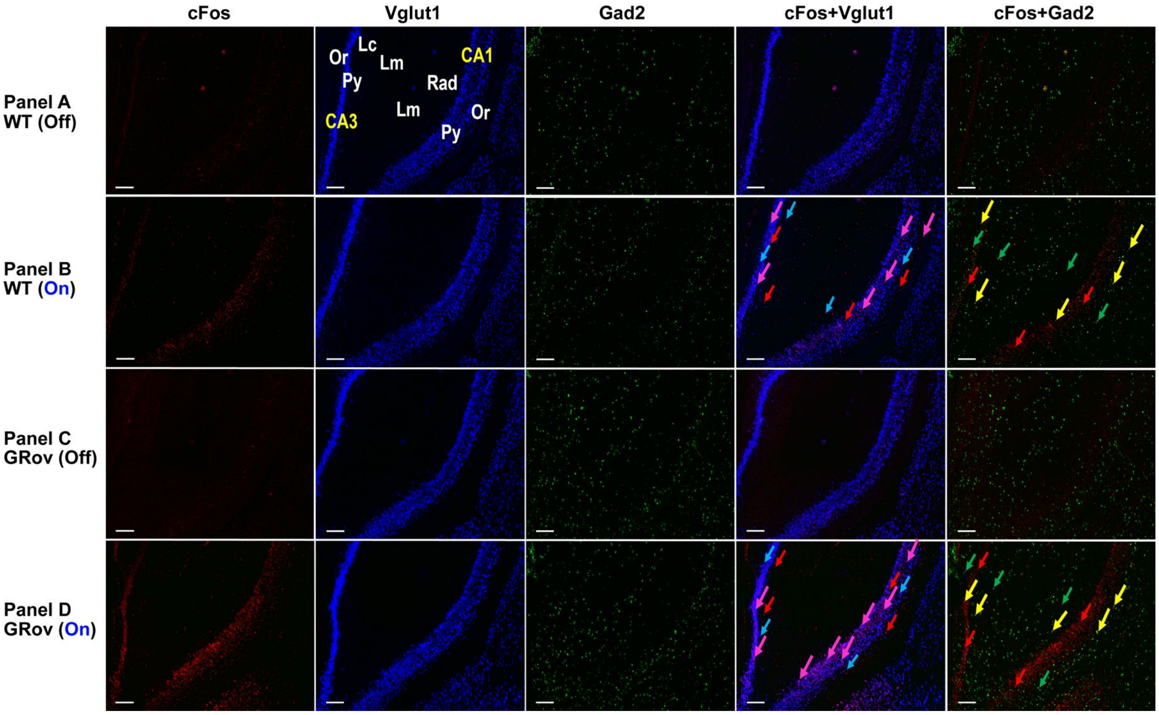


**Fig. S6.** Magnified views of CA3-CA1 of ventral hippocampus showing cFos colabeling with Vglut1$^+$ and Gad2$^+$ in the regions corresponding to the insets shown in Fig. S4. Predominant cFos+Vglut1 (Magenta) co-labeling was seen in the Py whereas cFos+Gad2 (Yellow) neurons were mostly localized in the Or, Rad, Lc and Lm layers of CA3. In the representative merged columns cFos+Vglut1 and cFos+Gad2 colocalizations are indicated by magenta and yellow arrows, whereas neurons expressing cFos, Vglut1 and Gad2 are indicated by red, blue, and green arrows, respectively. Scale bars, 100 μm.

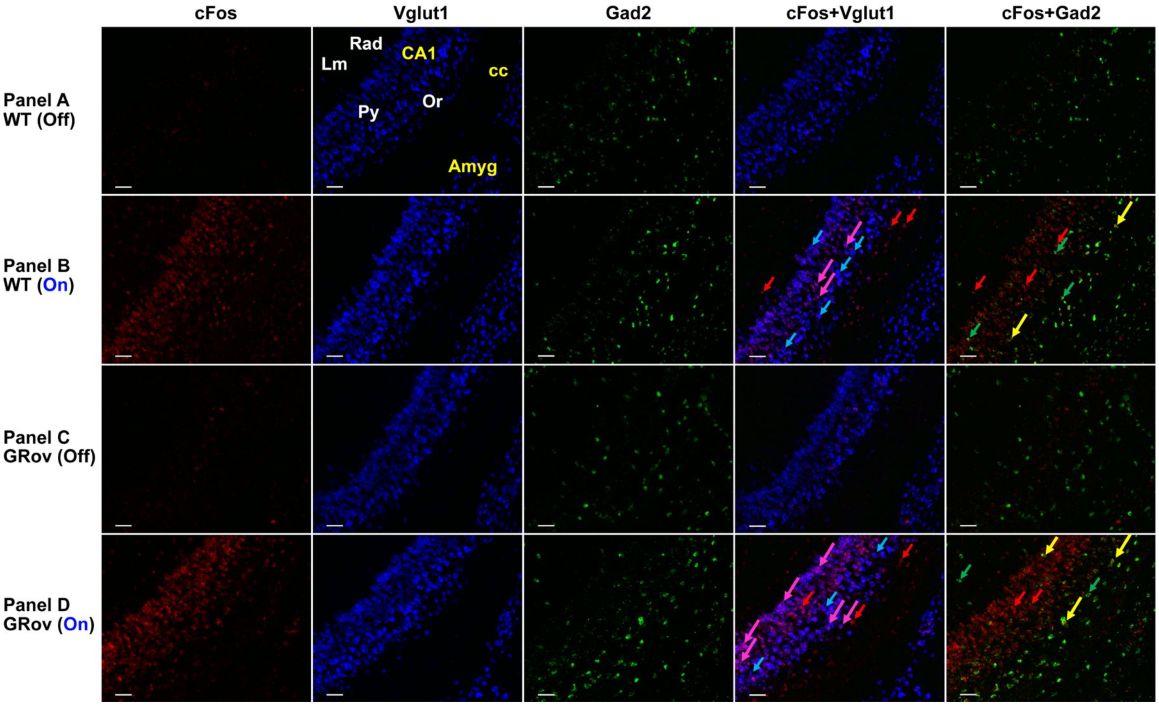


**Fig. S7.** Magnified views of CA1 of ventral hippocampus showing cFos co-labeling with Vglut1$^{+}$ and Gad2$^{+}$. Majority of the cFos+Vglut1 (Magenta) labeling can be seen in the Py whereas cFos+Gad2 (Yellow) neurons were mostly localized in the Or, Rad and Lm layers of CA1. In the representative merged columns cFos+Vglut1 and cFos+Gad2 colocalizations are indicated by magenta and yellow arrows, whereas neurons expressing cFos, Vglut1 and Gad2 are indicated by red, blue and green arrows, respectively. Scale bars, 50 μm.

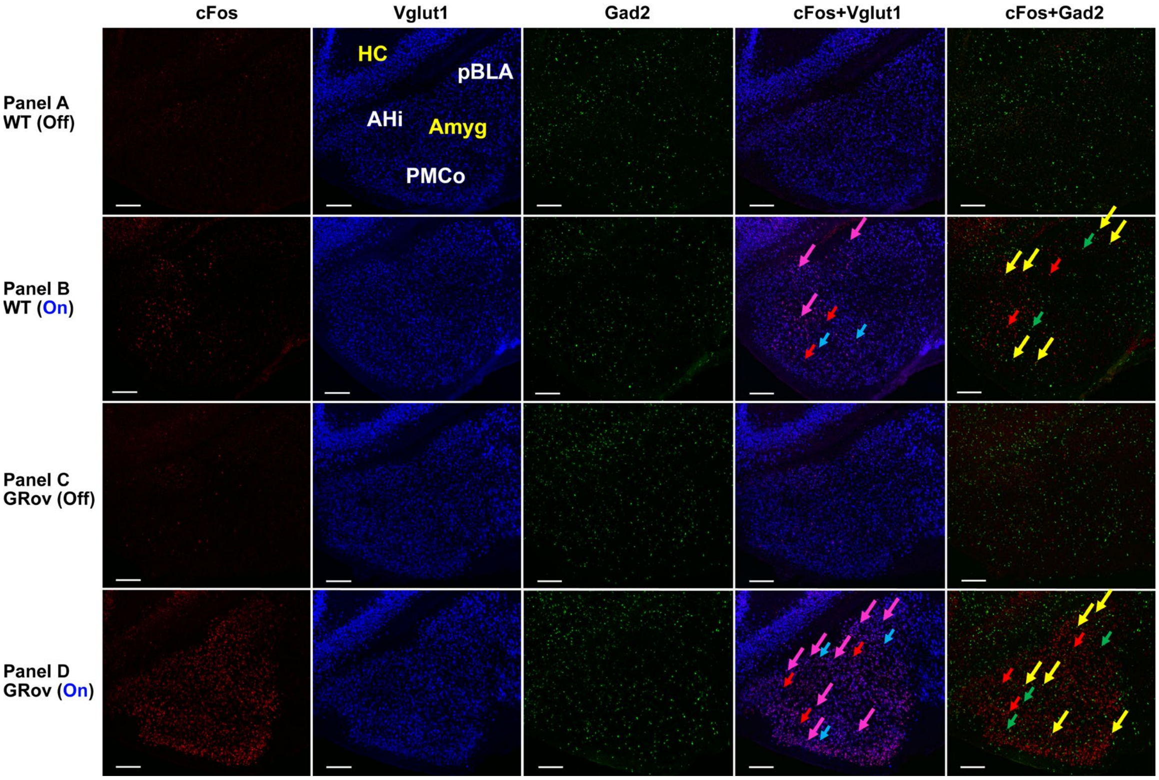


**Fig. S8.** Representative confocal images showing the increased cFos activity in the pBLA, AHi, and PMCo following light stimulation in vDG of WT and GRov mice (Panels B and D) compared to no stimulation groups (Panels A and C). Colocalization of cFos labeling (red) in Vglut1$^+$ neurons (blue) and Gad2$^+$ neurons (green) in the pBLA, AHi, and PMCo nuclei of amygdala illustrating predominant cFos activity in the glutamatergic neuronal population in the stimulated groups. Representative cFos+Vglut1 and cFos+Gad2 colocalization are indicated by magenta and yellow arrows, respectively and neurons expressing cFos, Vglut1 and Gad2 are indicated by red, blue and green arrows, respectively. Higher cFos activity co-localized with Vglut1$^+$ glutamatergic and Gad2$^+$ GABAergic neurons in the pBLA, AHi, and PMCo of both WT and GRov animals (Panels B, D) following the optogenetic stimulation in vDG. Scale bars, 200 μm.

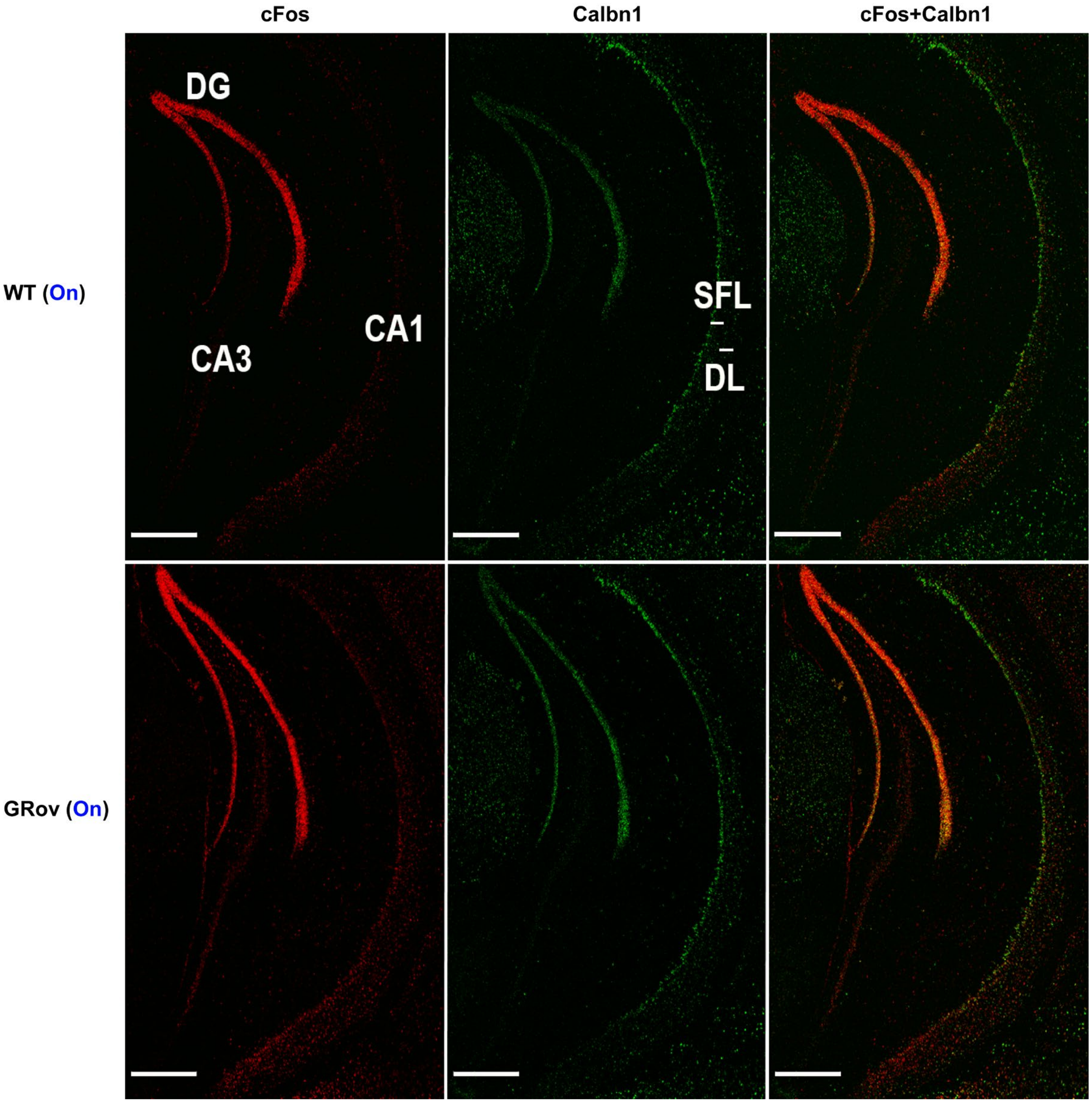


**Fig. S9.** Representative confocal photomicrographs showing increased cFos-based neuronal activation and its colocalization with Calbn1$^+$ expressing neurons in the DG and CA1 subregions of GRov-stimulated mice compared to WT-stimulated group following blue laser stimulation in vDG. HCR FISH labeling are shown with cFos (red), Calbn1 (green), cFos+calbn1 (yellow). Distinct distribution of Calbn1$^+$ neurons can be seen in the SFL of vCA1 compared to the DL. Compared to vDG and vCA1 subregions, vCA3 showed sparse and significantly low population of Calbn1$^+$ neurons. Scale bars, 500μm.

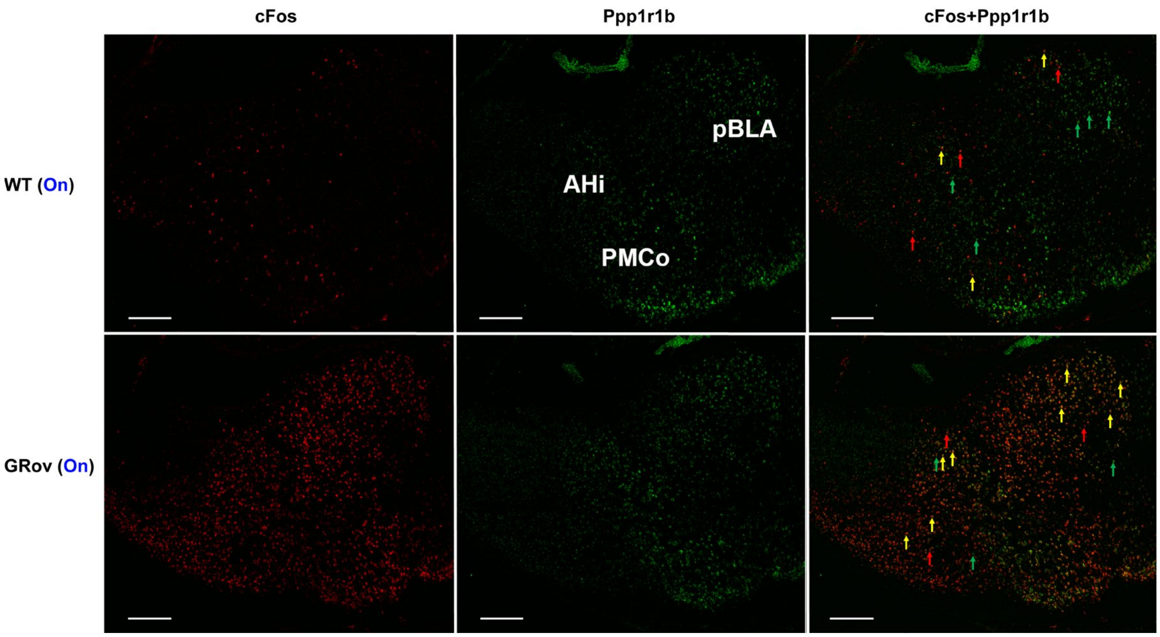


**Fig. S10.** Representative confocal photomicrographs of amygdaloid complex showing a greater colocalization (yellow) of cFos$^{+}$ neurons (red) with Ppp1r1b$^{+}$ expressing neurons (green) in pBLA, AHi, and PMCo subregions of GRov-stimulated mice compared to WT-stimulated group following blue laser stimulation in vDG. Representative cFos+Ppp1r1b colocalizations are indicated by yellow arrows, whereas neurons expressing cFos, and Ppp1r1b are indicated by red and green arrows, respectively. Scale bars, 250μm.

**Table S1. Distributions and relative abundance of cFos mRNA signal following optogenetic stimulation in dDG.**

| cFos positive regions | WT | | GRov | |
|---|---|---|---|---|
| | No stimulation | Stimulation | No stimulation | Stimulation |
| Orbital area | + | ++ | + | ++ |
| Olfactory bulb/nucleus/tubercle area | + | ++ | + | ++ |
| Piriform cortex, pyramidal layer | + | ++ | + | ++ |
| Anterior cingulate cortex | + | ++ | + | +++ |
| Prelimbic cortex | + | ++ | + | ++ |
| Infralimbic cortex | + | ++ | + | +++ |
| Nucleus accumbens | - | + | - | + |
| Bed nucleus of the stria terminalis | - | + | - | ++ |
| Lateral septal nucleus | + | ++++ | + | +++++ |
| Claustrum | + | ++ | + | ++ |
| Dorsal endopiriform nucleus | + | ++ | + | ++ |
| Primary/Secondary motor cortex, layer 1/2/3/6 | + | ++ | + | +++ |
| Primary somatosensory cortex, layer 1/2/3/6 | + | ++ | + | ++ |
| CA1 | + | +++ | + | ++++ |
| CA2 | + | ++ | + | ++ |
| CA3 | + | +++ | + | +++ |
| DG | + | +++++ | + | +++++ |
| Thalamus | + | ++ | + | ++ |
| Paraventricular hypothalamic nucleus | + | +++ | + | +++ |
| Zona incerta, anterior | - | ++ | - | ++ |
| Anterior hypothalamic nucleus | - | + | - | ++ |
| Periventricular hypothalamic nucleus | - | + | - | + |
| Ventromedial hypothalamic nucleus | - | + | - | + |
| Posterior hypothalamic nucleus | - | + | - | + |
| Arcuate nucleus, latero-posterior | - | + | - | ++ |
| Mammillary nucleus | + | ++ | + | ++ |
| Basolateral amygdala | + | ++ | + | ++ |
| Basolateral amygdaloid nucleus, posterior | + | ++ | + | +++ |
| Basomedial amygdaloid nucleus, posterior | - | + | - | + |
| Medial amygdaloid nucleus, posterior | + | ++ | + | ++ |
| Amygdalohippocampal area | + | ++ | + | +++ |
| Cortical amygdaloid area, posteromedial/posterolateral | + | ++ | + | +++ |
| Interpeduncular nucleus, caudal/lateral subnucleus | - | + | - | + |

The intensity of cFos mRNA signal was visually interpreted from ISH images and indicated as: -, undetectable; +, light; ++, moderate; +++, dense; ++++, heavy; +++++, compact. WT (light off), *n* = 5 mice; WT (light on), *n* = 7 mice; GRov (light off), *n* = 4 mice; GRov (light on), *n* = 7 mice.

**Table S2. Optical density for cFos mRNA in brain regions following optogenetic stimulation in dDG.**

| Regions | WT(Off) | WT(On) | GRov(Off) | GRov(On) |
|---|---|---|---|---|
| **M2** | 0.0397±0.0036 | 0.0666±0.0045*** | 0.037±0.0036 | 0.0619±0.0045** |
| **PFC** | 0.054±0.008 | 0.0777±0.0065* | 0.0560±0.013 | 0.0833±0.01 |
| **LS** | 0.0483±0.0058 | 0.1072±0.0061** | 0.0448±0.0064 | 0.1393±0.012**# |
| **PVN** | 0.0556±0.007 | 0.0808±0.0071* | 0.0492±0.0072 | 0.0912±0.0094** |
| **DG** | 0.0316±0.0023 | 0.4327±0.035*** | 0.037±0.0035 | 0.5847±0.046***# |
| **CA1** | 0.0298±0.0026 | 0.154±0.018*** | 0.0333±0.0014 | 0.2137±0.02***# |
| **CA3** | 0.0344±0.0045 | 0.1265±0.01*** | 0.0338±0.0027 | 0.1525±0.016*** |
| **pBLA** | 0.0286±0.0018 | 0.0395±0.0014*** | 0.0297±0.0025 | 0.0511±0.0051*# |
| **AHi** | 0.0308±0.0024 | 0.0465±0.0024*** | 0.0301±0.0019 | 0.0516±0.0026*** |
| **PMCo** | 0.0335±0.002 | 0.0472±0.0039*** | 0.0334±0.0015 | 0.0513±0.0016* |

Data expressed as mean ± SEM with WT (light off), $n$ = 5 mice for the no stimulation group; WT (light on), $n$ = 7 mice for the stimulation group; GRov (light off), $n$ = 4 mice for the no stimulation group; GRov (light on), $n$ = 7 mice for the stimulation group. Unpaired two-tailed $t$-test: *$P$ < 0.05, **$P$ < 0.01, ***P < 0.001 versus respective no stimulation group; $^{\#}P$ < 0.05 versus WT group under the same condition. M2, secondary motor cortex; PFC, prefrontal cortex; LS, lateral septum; PVN, paraventricular hypothalamic nucleus; DG, dentate gyrus; CA1, *cornu ammonis* field 1; CA3, *cornu ammonis* field 3; pBLA, posterior basolateral amygdala; AHi, amygdalohippocampal area; PMCo, posteromedial cortical amygdaloid nucleus.

**Table S3. Distributions and relative abundance of cFos mRNA signal following optogenetic stimulation in vDG.**

| cFos positive regions | WT | | GRov | |
|---|---|---|---|---|
| | No stimulation | Stimulation | No stimulation | Stimulation |
| Orbital area | + | ++ | + | ++ |
| Olfactory bulb/nucleus/tubercle area | + | ++ | + | ++ |
| Piriform cortex, pyramidal layer | + | ++ | + | ++ |
| Anterior cingulate cortex | + | ++ | + | ++ |
| Prelimbic cortex | + | ++ | + | ++ |
| Infralimbic cortex | + | ++ | + | ++ |
| Nucleus accumbens | - | + | - | ++ |
| Bed nucleus of the stria terminalis | - | + | - | ++ |
| Lateral septal nucleus | + | ++++ | + | +++++ |
| Claustrum | + | ++ | + | ++ |
| Dorsal endopiriform nucleus | + | ++ | + | ++ |
| Primary/Secondary motor cortex, layer 1/2/3/6 | + | ++ | + | ++ |
| Primary somatosensory cortex, layer 1/2/3/6 | + | ++ | + | ++ |
| CA1 | + | ++++ | + | +++++ |
| CA2 | + | ++ | + | ++ |
| CA3 | + | +++ | + | ++++ |
| DG | + | +++++ | + | +++++ |
| Thalamus | + | ++ | + | ++ |
| Paraventricular hypothalamic nucleus | + | +++ | + | +++ |
| Zona incerta, anterior | - | ++ | - | ++ |
| Anterior hypothalamic nucleus | - | + | - | + |
| Periventricular hypothalamic nucleus | + | ++ | + | ++ |
| Ventromedial hypothalamic nucleus | - | + | - | + |
| Posterior hypothalamic nucleus | + | ++ | + | ++ |
| Arcuate nucleus, latero-posterior | - | + | + | ++ |
| Mammillary nucleus | + | + | + | ++ |
| Basolateral amygdala | + | ++ | + | +++ |
| Basolateral amygdaloid nucleus, posterior | + | +++ | + | +++++ |
| Basomedial amygdaloid nucleus, posterior | + | ++ | + | +++ |
| Medial amygdaloid nucleus, posterior | + | ++ | + | ++ |
| Amygdalohippocampal area | + | +++ | + | +++++ |
| Cortical amygdaloid area, posteromedial/posterolateral | + | +++ | + | +++++ |
| Interpeduncular nucleus, caudal/lateral subnucleus | - | + | - | + |

The intensity of cFos mRNA signal was visually interpreted from ISH images and indicated as: -, undetectable; +, light; ++, moderate; +++, dense; ++++, heavy; +++++, compact. WT (light off), *n* = 4 mice; WT (light on), *n* = 5 mice; GRov (light off), *n* = 5 mice; GRov (light on), *n* = 5 mice.

**Table S4. Optical density for cFos mRNA in brain regions following optogenetic stimulation in vDG.**

| Regions | WT(Off) | WT(On) | GRov(Off) | GRov(On) |
|---|---|---|---|---|
| **M2** | 0.0411±0.0023 | 0.0526±0.0043* | 0.0358±0.0025 | 0.0946±0.0235* |
| **PFC** | 0.0549±0.0056 | 0.0716±0.011 | 0.0584±0.011 | 0.1233±0.0345 |
| **LS** | 0.0452±0.0035 | 0.1269±0.0125** | 0.0443±0.0046 | 0.1626±0.0173** |
| **PVN** | 0.0563±0.0064 | 0.0906±0.0154 | 0.0448±0.0028 | 0.1124±0.011*** |
| **DG** | 0.0285±0.0031 | 0.4999±0.042*** | 0.029±0.0015 | 0.626±0.0151***# |
| **CA1** | 0.027±0.0017 | 0.1617±0.0235*** | 0.0279±0.0012 | 0.2572±0.0249***# |
| **CA3** | 0.03±0.0023 | 0.1107±0.0119*** | 0.0293±0.0015 | 0.1651±0.0237*** |
| **pBLA** | 0.0266±0.0019 | 0.086±0.0289 | 0.0258±0.0007 | 0.2311±0.0525**# |
| **AHi** | 0.0305±0.0028 | 0.0901±0.0242* | 0.0293±0.0013 | 0.2792±0.0733**# |
| **PMCo** | 0.0317±0.0017 | 0.118±0.0343* | 0.0327±0.0014 | 0.3561±0.0868**# |

Data expressed as mean ± SEM with WT (light off), $n$ = 4 mice for the no stimulation group; WT (light on), $n$ = 5 mice for the stimulation group; GRov (light off), $n$ = 5 mice for the no stimulation group; GRov (light on), $n$ = 5 mice for the stimulation group. Unpaired two-tailed $t$-test: *$P < 0.05$, **$P < 0.01$, ***$P < 0.001$ versus respective no stimulation group; $^{\#}P < 0.05$ versus WT group under the same condition.

**Table S5. Estimated number density of cFos-colocalized glutamatergic and GABAergic neurons in the hippocampal formation following optogenetic stimulation of vDG.**

| Group | Cell type | vDG | vCA1 | vCA3 |
|---|---|---|---|---|
| WT (Off) | ***cFos***$^{+}$ | 18.66±3.9 | 21.54±2.97 | 22.88±1.86 |
| | **cFos**$^{+}$**Vglut1**$^{+}$ | 10.62±1.45 | 12.21±2.61 | 14.45±1.56 |
| | **Vglut1**$^{+}$ | 764.55±59.69 | 542.25±28.09 | 627.73±69.06 |
| | **cFos**$^{+}$**Gad2**$^{+}$ | 5.93±1.8 | 7.7±1.32 | 5.16±1.32 |
| | **Gad2**$^{+}$ | 86.32±10.73 | 89.36±23.7 | 87.86±16.18 |
| WT (On) | **cFos**$^{+}$ | 731.89±52.27*** | 224.4±20.51*** | 184.77±24.51** |
| | **cFos**$^{+}$**Vglut1**$^{+}$ | 684.04±46.93*** | 136.26±24.34** | 120.06±11** |
| | **Vglut1**$^{+}$ | 725.76±60.61 | 461.26±71.16 | 561.81±60.69 |
| | **cFos**$^{+}$**Gad2**$^{+}$ | 46.95±10.85*** | 58.96±9.05** | 54.42±17.32*** |
| | **Gad2**$^{+}$ | 58.9±22.23 | 83.6±10.73 | 79.62±24.55 |
| GRov (Off) | **cFos**$^{+}$ | 12.66±1.97 | 23.87±2.71 | 21.55±2.53 |
| | **cFos**$^{+}$**Vglut1**$^{+}$ | 5.03±0.79 | 7.1±1.7 | 11.43±1.28 |
| | **Vglut1**$^{+}$ | 734.78±76.75 | 548.24±83.88 | 588.29±33.11 |
| | **cFos**$^{+}$**Gad2**$^{+}$ | 6.39±2.11 | 11.67±4.29 | 7.27±1.85 |
| | **Gad2**$^{+}$ | 78.32±15.85 | 97.83±24.97 | 91.98±38.97 |
| GRov (On) | **cFos**$^{+}$ | 837.42±51.13*** | 345.74±20.4***## | 270.68±23*** |
| | **cFos**$^{+}$**Vglut1**$^{+}$ | 773.2±48.81*** | 231.58±8.32***# | 191.84±34.73** |
| | **Vglut1**$^{+}$ | 808.07±63.52 | 554.29±58.98 | 605.56±60.57 |
| | **cFos**$^{+}$**Gad2**$^{+}$ | 56.62±12.64*** | 43.16±9.88***# | 52.95±18.2*** |
| | **Gad2**$^{+}$ | 76.64±25.95 | 74.62±17.53 | 84.14±26.14 |

Data expressed as mean ± SEM per 2.5X10$^{6}$ μm$^{2}$ area with WT (light off), $n$ = 3 mice for the no stimulation group; WT (light on), $n$ = 4 mice for the stimulation group; GRov (light off), $n$ = 3 mice for the no stimulation group; GRov (light on), $n$ = 5 mice for the stimulation group. Significant comparisons between control and stimulated groups are depicted with $^{**}P < 0.01$, $^{***}P < 0.001$. Genotype-based significant comparisons between WT- and GRov-stimulated groups are shown as $^{\#}P < 0.05$, $^{\#\#}P < 0.01$.

**Table S6. Estimated number density of cFos-colocalized glutamatergic and GABAergic neurons in the amygdaloid complex following optogenetic stimulation of vDG.**

| Group | Cell Type | pBLA | AHi | PMCo |
|---|---|---|---|---|
| **WT (Off)** | **cFos$^+$** | 32.97±5.76 | 26.05±3.98 | 18.39±1.79 |
| | **cFos$^+$Vglut1$^+$** | 22.39±3.89 | 17.29±2.29 | 11.91±1.28 |
| | **Vglut1$^+$** | 604.82±76.21 | 790.87±74.03 | 643.52±64.77 |
| | **cFos$^+$Gad2$^+$** | 4.06±1.65 | 5.82±2.54 | 3.06±0.78 |
| | **Gad2$^+$** | 67.14±15.32 | 116.74±25.6 | 73.08±12.64 |
| **WT (On)** | **cFos$^+$** | 153.26±48.93** | 219.16±33.09*** | 191.54±44.8*** |
| | **cFos$^+$Vglut1$^+$** | 81.8±8.16** | 157.36±20.5** | 154.31±14.08** |
| | **Vglut1$^+$** | 633.81±41.33 | 795.64±87.08 | 657.86±66.52 |
| | **cFos$^+$Gad2$^+$** | 24.51±12.87* | 22.54±5.58** | 12.32±3.65** |
| | **Gad2$^+$** | 106.77±48.98 | 112.55±42.89 | 81.58±28.82 |
| **GRov (Off)** | **cFos$^+$** | 33.29±2.19 | 32.02±0.49 | 18.41±3.76 |
| | **cFos$^+$Vglut1$^+$** | 25.1±1.67 | 25.14±0.39 | 12.75±2.44 |
| | **Vglut1$^+$** | 590.53±34.38 | 901.88±77.12 | 728.99±20.09 |
| | **cFos$^+$Gad2$^+$** | 4.84±2.06 | 6±2.77 | 3.21±0.88 |
| | **Gad2$^+$** | 83.75±20.36 | 138.68±11.02 | 63.36±16.99 |
| **GRov (On)** | **cFos$^+$** | 318.23±80.72***## | 527.41±98.37***### | 411.64±79.05***## |
| | **cFos$^+$Vglut1$^+$** | 320.9±35.76**## | 470.55±60.08***## | 345.75±45.88**## |
| | **Vglut1$^+$** | 630.31±58.4 | 847.54±86.17 | 641.23±77.24 |
| | **cFos$^+$Gad2$^+$** | 19.45±4.73** | 26.87±4.75*** | 10.2±2.35** |
| | **Gad2$^+$** | 95.74±25.46 | 109.57±28.49 | 57.63±11.2 |

Data expressed as mean ± SEM per $2.5X10^6$ $\mu m^2$ area with WT (light off), $n$ = 3 mice for the no stimulation group; WT (light on), $n$ = 4 mice for the stimulation group; GRov (light off), $n$ = 3 mice for the no stimulation group; GRov (light on), $n$ = 5 mice for the stimulation group. Significant comparisons between control and stimulated groups are depicted with *$P$ < 0.05, **$P$ < 0.01, ***$P$ < 0.001. Genotype-based significant comparisons between WT- and GRov-stimulated groups are shown as $^{\#\#}P$ < 0.01, $^{\#\#\#}P$ < 0.001.

**Table S7. Estimated number density of cFos-colocalized Calbn1 in the hippocampal formation following optogenetic stimulation of vDG.**

| Group | Cell Type | DG | CA1 | CA3 |
|---|---|---|---|---|
| WT (On) | **cFos⁺Calbn1⁺** | 291.88±33.71 | 81.03±10.16 | 7.4±1.32 |
| | **Calbn1⁺** | 365.88±26.97 | 230.75±24.02 | 16.08±1.31 |
| GRov (On) | **cFos⁺Calbn1⁺** | 287.67±28.25 | 143.42±14.28$^{\#}$ | 8.1±1.38 |
| | **Calbn1⁺** | 378.02±30.28 | 228±19.49 | 15.11±2.06 |

Data expressed as mean ± SEM per 2.5X10$^6$ μm$^2$ area with WT (light on), *n* = 4 mice; GRov (light on), *n* = 5 mice. Genotype-based significant comparisons are depicted with $^{\#}P < 0.05$.

**Table S8. Estimated number density of cFos-colocalized Ppp1r1b neurons in the amygdaloid complex following optogenetic stimulation of vDG.**

| Group | Cell Type | pBLA | AHi | PMCo |
|---|---|---|---|---|
| WT (On) | **cFos⁺Ppp1r1b⁺** | 35.36±4.88 | 43.34±9.72 | 27.46±4.42 |
| | **Ppp1r1b⁺** | 200.76±12.7 | 137.7±10.95 | 130.07±13.3 |
| GRov (On) | **cFos⁺Ppp1r1b⁺** | 152.56±16.48$^{\#\#\#}$ | 75.51±8.82$^{\#}$ | 65.45±5.96$^{\#\#}$ |
| | **Ppp1r1b⁺** | 191.71±10.13 | 133.77±10.45 | 127.42±8.16 |

Results are expressed as mean ± SEM per 2.5X10$^6$ μm$^2$ area with WT (light on), *n* = 4 mice; GRov (light on), *n* = 5 mice. Genotype-based significant comparisons are depicted with $^{\#}P < 0.05$, $^{\#\#}P < 0.01$, $^{\#\#\#}P < 0.001$.

### Table S9. List of split-initiator DNA probe (HCR version 2.0) sequences.

#### **S9a. cFos Probe:** NM_010234.3; HCR amplifier: B1-AlexaFluor647

| Odd | 1st half of Initiator I1 + Spacer (AA) + Probe Sequence | Even | Probe Sequence + Spacer (TA) + 2nd half of Initiator I1 |
|---|---|---|---|
| 1 | gAggAgggCAgCAAACggAAGAGACACAGACCAGGCCTTGACTCA | 2 | ATGAGAAAACTAAGGAGAAAGAGAATAgAAgAgTCTTCCTTTACg |
| 3 | gAggAgggCAgCAAACggAATGAACCCAACAGATTAGTTAGTGCT | 4 | AATCCAGCACCAGGTTAATTCCAATTAgAAgAgTCTTCCTTTACg |
| 5 | gAggAgggCAgCAAACggAATGTTAAAATCAGCTGCACTAGATAC | 6 | CGCTATTGCCAGGAACACAGTAGGTTAgAAgAgTCTTCCTTTACg |
| 7 | gAggAgggCAgCAAACggAAATATTGGTCGTTTCTAATTGGAACA | 8 | AATAAAGTCCTATCTTTTCTTAGTTTAgAAgAgTCTTCCTTTACg |
| 9 | gAggAgggCAgCAAACggAAAGCTATTGATTTCTATCTACTGGAA | 10 | CGCTGAAGGACTACAGTACATGGATTAgAAgAgTCTTCCTTTACg |
| 11 | gAggAgggCAgCAAACggAATCAGTAACATGACAATGAACATTGA | 12 | CATTCAGACCACCTCGACAATGCATTAgAAgAgTCTTCCTTTACg |
| 13 | gAggAgggCAgCAAACggAATCATGGAAAACTGTTAATGTCAGAA | 14 | TAAATTGAAAACACAATAAAAACGTTAgAAgAgTCTTCCTTTACg |
| 15 | gAggAgggCAgCAAACggAAATATCTGAGAATCCATCTTAATAAA | 16 | GTAGAAAAAAATAAAATAAAAATATTAgAAgAgTCTTCCTTTACg |
| 17 | gAggAgggCAgCAAACggAATTTCCACATGTCGAAAGACCTCAGG | 18 | TGCTTAAAATTTTTCATTCAAATTCTAgAAgAgTCTTCCTTTACg |
| 19 | gAggAgggCAgCAAACggAAATGTCTTGGAACAATAAGCAAACAA | 20 | CATTCAACTTAAATGCTTTTATTGATAgAAgAgTCTTCCTTTACg |

#### **S9b. Vglut1 Probe:** NM_182993.2; HCR amplifier: B3-AlexaFluor594

| Odd | 1st half of Initiator I1 + Spacer (TT) + Probe Sequence | Even | Probe Sequence + Spacer (TT) + 2nd half of Initiator I1 |
|---|---|---|---|
| 1 | gTCCCTgCCTCTATATCTTTACTGCCCCACAGTGGGAGGCCCGTG | 2 | GAGGTGTATGGAGTGGAAGTCCTGGTTCCACTCAACTTTAACCCg |
| 3 | gTCCCTgCCTCTATATCTTTTGGGAACAAGGGAGGACTTGCATCT | 4 | AGGGAAAGAGGGCTGGTCGGACAGCTTCCACTCAACTTTAACCCg |
| 5 | gTCCCTgCCTCTATATCTTTTCACCCCCCGCAGGAGGCAGTTGAG | 6 | TATCCTTGAAACTGCTAGTGTGCAGTTCCACTCAACTTTAACCCg |
| 7 | gTCCCTgCCTCTATATCTTTTTAGGCGAGCCTTGAAACTAATAGA | 8 | AACCAGCTTGGAAAAATGTAGAATTTTCCACTCAACTTTAACCCg |
| 9 | gTCCCTgCCTCTATATCTTTAACGGCGGCATTGGTGGTTAGGTTA | 10 | GAAACGCTGGTGAGAATCAGTCCTGTTCCACTCAACTTTAACCCg |
| 11 | gTCCCTgCCTCTATATCTTTGCCCGGCAAAGTGTGCTGGTGAGGG | 12 | CAATGATTGTACTAAGCTAAGGTCATTCCACTCAACTTTAACCCg |
| 13 | gTCCCTgCCTCTATATCTTTAGCCACTACTGAGACCTGAAAACTG | 14 | CGAGCCGCTGAATTAATAGCTTTGGTTCCACTCAACTTTAACCCg |
| 15 | gTCCCTgCCTCTATATCTTTGACACACAACAAATGGCCACTGAGA | 16 | AGATTTGGAATCATTTAGCCCCTGATTCCACTCAACTTTAACCCg |
| 17 | gTCCCTgCCTCTATATCTTTTGTAACTTCTCTCACACACCTCACC | 18 | CCAGCCCCGCTCCCTTTCCTGGGATTTCCACTCAACTTTAACCCg |

#### **S9c. Gad2 Probe:** NM_008078.2; HCR amplifier: B2-AlexaFluor546

| Odd | 1st half of Initiator I1 + Spacer (AA) + Probe Sequence | Even | Probe Sequence + Spacer (AA) + 2nd half of Initiator I1 |
|---|---|---|---|
| 1 | CCTCgTAAATCCTCATCAAAAGGGTAGAAGAGAGGCAAGGACAGG | 2 | CACAGCTTGGGACTGGGTGAAAGGGAAATCATCCAgTAAACCgCC |
| 3 | CCTCgTAAATCCTCATCAAATCTCTAAGAGCCAATGGAGAGGGCA | 4 | GGGTGGGACTTAGTTGAGGTTATGTAAATCATCCAgTAAACCgCC |
| 5 | CCTCgTAAATCCTCATCAAAACACAGTTGTCAAAGAGATTTCTTA | 6 | CAGAGATGAAGCATTTTGTGTGCCAAAATCATCCAgTAAACCgCC |
| 7 | CCTCgTAAATCCTCATCAAAGCTCCATAATTGGTTTCTCTGGCCT | 8 | ATACCTGCACTGTCAGCAGCCTGTGAAATCATCCAgTAAACCgCC |
| 9 | CCTCgTAAATCCTCATCAAACCTTCTCCAAGACCCTGTAGAGTCA | 10 | CCTTTGTCCATGTTCTGAGGAGCAGAAATCATCCAgTAAACCgCC |
| 11 | CCTCgTAAATCCTCATCAAATGTTACTATATTTACACCTGTGCAT | 12 | TGGTTTGATGTTTTTGCTTCTTTGTAAATCATCCAgTAAACCgCC |
| 13 | CCTCgTAAATCCTCATCAAAAGAAAAGCACGTGCAAGATGATACC | 14 | CAGTGTTTGCATTTAGCACTTGAAAAAATCATCCAgTAAACCgCC |
| 15 | CCTCgTAAATCCTCATCAAATTTGCATACACAACTAATTAATACA | 16 | TCAGAAACACCATTTGGCAACAAGAAAATCATCCAgTAAACCgCC |
| 17 | CCTCgTAAATCCTCATCAAAAGGTTGCCACATTTGTTTTTATTTT | 18 | CATACACAAGTTTATATTTGGTAGCAAATCATCCAgTAAACCgCC |
| 19 | CCTCgTAAATCCTCATCAAAGAAAACACAGGTAAAATACTTTGAT | 20 | ATTTACACATTTATTTGGGTTTAGAAAATCATCCAgTAAACCgCC |

### **S9d. Calbn1 Probe:** NM_009788.4; HCR amplifier: B4-AlexaFluor546

| Odd | 1st half of Initiator I1 + Spacer (AA) + Probe Sequence | Even | Probe Sequence + Spacer (AA) + 2nd half of Initiator I1 |
|---|---|---|---|
| 1 | CCTCAACCTACCTCCAACAACGCAGGCTGAGCTGGGCGCGGCGCG | 2 | GAGCGCGAACCTCGGGCGAGAGGGCATTCTCACCATATTCgCTTC |
| 3 | CCTCAACCTACCTCCAACAACCTTCCAGGTAACCACTTCCGTCAG | 4 | TCCTGGATCAAGTTCTGCAGCTCCTATTCTCACCATATTCgCTTC |
| 5 | CCTCAACCTACCTCCAACAATCTGTGGGTAAGACGTGAGCCAACT | 6 | CATCGAAAGAGCAGCAAGAAATTCTATTCTCACCATATTCgCTTC |
| 7 | CCTCAACCTACCTCCAACAAGTCTTGTTTGCTTTCTCTAGTAGGT | 8 | TACTCTGCTAGTTTTGTATCATCCAATTCTCACCATATTCgCTTC |
| 9 | CCTCAACCTACCTCCAACAATTCCCACACATTTTGATTCCCTGGA | 10 | TATAACTCAAAAGCCTTATTGAACTATTCTCACCATATTCgCTTC |
| 11 | CCTCAACCTACCTCCAACAAAAGGCCATTATGTTCTTCTTGTATG | 12 | GTTCGGTACAGCTTCCCTCCATCCGATTCTCACCATATTCgCTTC |
| 13 | CCTCAACCTACCTCCAACAAGATATAAAAGAAAATACAGCCTACT | 14 | AGTTCTCTATATGCAGTAGAATTTAATTCTCACCATATTCgCTTC |
| 15 | CCTCAACCTACCTCCAACAATTTTTCAAAATTGATTGTACTGACT | 16 | TATAAACGCAAAACCATGGTAGATTATTCTCACCATATTCgCTTC |
| 17 | CCTCAACCTACCTCCAACAAGCCTAAAATAAAAACAGTTGGCTTG | 18 | GGTGCTTTGGGTGACAGTCCTATGGATTCTCACCATATTCgCTTC |
| 19 | CCTCAACCTACCTCCAACAAAGACATGTCACTGATCAGTACACTA | 20 | CAATAGTGTGGTTTATATTTAAGCAATTCTCACCATATTCgCTTC |

### **S9e. Ppp1r1b Probe:** NM_001313970.1; HCR amplifier: B2-AlexaFluor594

| Odd | 1st half of Initiator I1 + Spacer (TT) + Probe Sequence | Even | Probe Sequence + Spacer (TT) + 2nd half of Initiator I1 |
|---|---|---|---|
| 1 | CCTCgTAAATCCTCATCAAACCACCCCCCCTCGGGACGTGGACTT | 2 | CCGGGACTCTGGCTTTAACCTCTTCAAATCATCCAgTAAACCgCC |
| 3 | CCTCgTAAATCCTCATCAAAGGGTGCGGAGTGGGAGCCAGGGCTA | 4 | GAAGCGTGTTCTCCCGGGCTGGGAGAAATCATCCAgTAAACCgCC |
| 5 | CCTCgTAAATCCTCATCAAAGCGCCTCTGCCGCTACTGGCTGCAC | 6 | ATCCCCCACTCTGCCTTCTTCCTCTAAATCATCCAgTAAACCgCC |
| 7 | CCTCgTAAATCCTCATCAAAGGCCTTGCTCTACCAGCTTCTGTCT | 8 | CCTGAGCTATCTCGGCGGGCTGCCCAAATCATCCAgTAAACCgCC |
| 9 | CCTCgTAAATCCTCATCAAAGTCCCCAGGTCAGAGCTCAGGATCC | 10 | GGGCTCGCCTGCAGCGCCGGGCTCAAAATCATCCAgTAAACCgCC |
| 11 | CCTCgTAAATCCTCATCAAACGACGGTCGGGAGGACTGAACAGCG | 12 | AATCCCGCGCTCCACGAGGAGAACGAAATCATCCAgTAAACCgCC |
| 13 | CCTCgTAAATCCTCATCAAAGGAGCACTGTCCCCGCACCCAGGAA | 14 | TGGCGCGTGCTGCGCGGAGGAGGAGAAATCATCCAgTAAACCgCC |
| 15 | CCTCgTAAATCCTCATCAAATTCTCTGATGTGGAGAGGCCTCCTC | 16 | ACTTTGGGTGGTGCCCCTCTCCAGAAAATCATCCAgTAAACCgCC |
| 17 | CCTCgTAAATCCTCATCAAACAAGTTGCTAATGGTCTGCAGGTGC | 18 | CTCTTCCTCCGAGGCCTGGTTCTCAAAATCATCCAgTAAACCgCC |
| 19 | CCTCgTAAATCCTCATCAAAGCTCCCGAAGCTCCCCTAACTCATC | 20 | CCTCATCATCCTCCTGTGGGTACCCAAATCATCCAgTAAACCgCC |

Listed mouse (*Mus musculus*) probes were designed in lab and synthesized-purchased from IDT. Sequences are listed 5' to 3'.
